# Interpretable biomarker discovery from small-sample microarray datasets using XGBoost rank aggregation and SVM-RFECV

**DOI:** 10.64898/2026.08.13.744652

**Authors:** Mashiyat Mahjabin Prapty, Mohammad Saifur Rahman

## Abstract

**Motivation:** High-dimensional microarray datasets remain valuable for cancer biomarker discovery, but their small sample sizes make robust and interpretable feature selection challenging. Efficient workflows are needed to derive compact gene signatures while preserving biological interpretability.

**Results:** We developed a two-stage biomarker-discovery workflow that combines cross-validated XGBoost rank aggregation with support vector machine recursive feature elimination and cross-validation (SVM-RFECV) to identify compact candidate biomarker panels. The workflow was evaluated on 21 public binary and multiclass microarray datasets using repeated stratified cross-validation for internal validation. Across the dataset collection, the selected panels demonstrated strong internal discriminative performance while remaining sufficiently compact for downstream biological interpretation. SHAP analysis identified dataset- and class-specific discriminative genes, and functional enrichment analysis supported the biological coherence of representative consensus signatures. The proposed workflow provides an interpretable and reproducible framework for candidate biomarker discovery from small-sample microarray datasets.

**Availability:** Source code and processed outputs are freely available at https://github.com/mashiyat-mahjabin-prapty/microarray-feature-selection.

## Introduction

Cancer remains one of the leading causes of morbidity and mortality worldwide, making the identification of reliable molecular biomarkers important for diagnosis, prognosis, subtype classification, and personalized treatment. DNA microarray technology enabled the simultaneous measurement of thousands of gene-expression levels and established an important foundation for computational biomarker discovery. Since the landmark study by Golub et al. (1999) showed that acute myeloid leukemia (AML) and acute lymphoblastic leukemia (ALL) could be distinguished from gene-expression profiles, microarray analysis has remained an important paradigm for molecular cancer classification and biomarker discovery (Golub et al., 1999). Although RNA sequencing is now widely used, public microarray datasets remain valuable for benchmarking biomarker-discovery workflows because many well-studied cancer classification datasets are publicly available and continue to be used in feature-selection research (Wang et al., 2025).

A defining characteristic of microarray datasets is their small-sample, high-dimensional structure, where thousands to tens of thousands of measured probes are available for a limited number of biological samples. Such settings increase the risk of overfitting, reduce model generalizability, increase computational complexity, and make feature selection sensitive to sampling variation (Hambali et al., 2020; Momeni et al., 2020). Consequently, feature selection is an essential component of microarray analysis, not only for improving predictive performance but also for identifying compact and biologically meaningful candidate biomarker panels. Compared with feature extraction, feature selection preserves the original gene representation, thereby maintaining biological interpretability while reducing computational burden and model complexity (Wang et al., 2025).

Feature-selection methods are commonly categorized into filter, wrapper, embedded, and hybrid approaches (Chandrashekar and Sahin, 2014; Wang et al., 2025). Filter methods rank features independently of the learning algorithm using statistical or information-theoretic criteria, providing computational efficiency but often failing to capture gene-gene interactions. Wrapper methods evaluate candidate subsets using predictive models and can achieve strong predictive performance, but they are computationally expensive when applied directly to tens of thousands of probes. Embedded methods integrate feature selection into model training, whereas hybrid methods combine rapid preliminary screening with classifier-guided refinement. Because microarray datasets typically contain extremely high-dimensional feature spaces but relatively few samples, hybrid strategies are attractive because they reduce the search space before more focused wrapper refinement is performed (Got et al., 2021).

Recent studies have proposed increasingly sophisticated feature-selection frameworks for microarray classification. Deterministic hybrid methods combine filtering and wrapper refinement to improve predictive performance while reducing computational cost (Li et al., 2023). Evolutionary and swarm-intelligence approaches, including AltWOA, MGWO, MUL-MGO, and MMODE, explore the large combinatorial search space of candidate gene subsets (Kundu et al., 2022; Pan et al., 2023; Osama et al., 2023; Li et al., 2024). Other recent directions include graph-based learning, Bayesian ensemble feature selection, class-imbalance-aware scoring, and ensemble similarity-based methods designed to improve feature-selection stability and predictive performance (Jenul et al., 2022; Xie et al., 2024; Roy et al., 2024; Khan et al., 2024). Collectively, these studies demonstrate substantial progress, but they also highlight recurring challenges: computationally intensive optimization, extensive hyperparameter tuning, heterogeneous validation protocols, and uneven downstream biological interpretation.

Despite these advances, relatively few studies present an end-to-end biomarker-discovery workflow that jointly emphasizes stable feature screening, reproducible feature refinement, model interpretability, and downstream biological contextualization. Many published methods primarily focus on optimizing feature-selection algorithms or maximizing classification accuracy, whereas biological interpretation is often limited to reporting selected genes without systematically examining their contribution to model predictions or their functional relevance. Furthermore, heterogeneous preprocessing pipelines, classifiers, selected-gene reporting practices, and validation protocols make direct comparison among published studies inherently difficult. These observations motivate practical and reproducible biomarker-discovery workflows that integrate computational feature selection with explainable machine learning and functional biological interpretation rather than treating these components independently (Hambali et al., 2020; Abd-Elnaby et al., 2021; Wang et al., 2025).

To address these challenges, we present an interpretable two-stage biomarker-discovery workflow for small-sample cancer microarray datasets. The proposed framework first applies cross-validated XGBoost rank aggregation to identify recurrent high-ranking candidate features, followed by support vector machine recursive feature elimination with cross-validation (SVM-RFECV) to generate compact candidate biomarker panels. XGBoost provides an efficient embedded feature-ranking mechanism capable of handling high-dimensional data (Chen and Guestrin, 2016), while RFECV performs classifier-guided wrapper refinement to eliminate redundant features and identify compact predictive gene sets (Guyon et al., 2002). The resulting panels are evaluated using repeated internal cross-validation and interpreted through SHapley Additive exPlanations (SHAP) (Lundberg and Lee, 2017) together with functional enrichment analysis. Rather than introducing a new optimization algorithm, our objective is to develop a practical, reproducible, and biologically interpretable workflow that integrates feature screening, deterministic wrapper refinement, internal validation, explainable machine learning, and downstream biological analysis within a unified framework. The workflow is evaluated across 21 publicly available binary and multiclass microarray datasets, representing diverse cancer classification tasks.

The principal contributions of this study are summarized as follows:

1. We propose an interpretable two-stage biomarker-discovery workflow that combines cross-validated XGBoost feature-rank aggregation with deterministic SVM-RFECV refinement to generate compact candidate biomarker panels.
2. We evaluate the proposed workflow across 21 publicly available binary and multiclass microarray datasets using repeated stratified internal validation and adaptive cross-validation strategies appropriate for small-sample datasets.
3. We report consensus biomarker panels, fold-level feature-selection frequencies, and feature-selection stability summaries to facilitate reproducible interpretation of selected genes.
4. We integrate SHAP-based feature attribution with functional enrichment analysis to connect computationally selected biomarkers with their potential biological relevance.

## Materials and Methods

### Datasets

Twenty-one classification tasks derived from publicly available disease datasets, comprising both binary and multiclass classification problems. The datasets varied substantially in sample size and original dimensionality, providing a heterogeneous benchmark for evaluating the proposed biomarker-discovery workflow under small-sample, high-dimensional conditions. Dataset names, numbers of samples, original feature counts, class distributions, and internal-validation results are reported in Table 2. The analyses were performed using the expression matrices and class labels provided with the processed datasets.

The datasets represented three broad prediction settings: tumor versus non-malignant classification, disease or tumor subtype classification, and relapse prediction. Sample sizes ranged from 36 to 253, while the number of measured features ranged from 1,626 to 54,613. Because several datasets contained small minority classes, the number of cross-validation folds was adapted to the smallest class size when required.

### Preprocessing

Each dataset was loaded as a tabular expression matrix with samples as rows, probe-level measurements as features, and a class-label column. Probe-level features were retained during model fitting so that multiple probes mapping to the same gene were not prematurely collapsed. Missing values were handled using mean imputation fitted only within the relevant training fold or model-fitting procedure. Standardization was applied within scikit-learn pipelines for scale-sensitive models, including logistic regression, support vector machines, and the linear SVM estimator used by RFECV. Thus, imputation and scaling were performed inside the corresponding cross-validation procedures rather than globally before model fitting.

### Feature Selection

Feature selection was performed in two stages. Cross-validated XGBoost ranking first reduced the original high-dimensional expression matrix to a candidate set of 500 probes. These candidates were subsequently refined using support vector machine recursive feature elimination with cross-validation (SVM-RFECV). This combination provided efficient initial screening followed by classifier-guided selection of a compact biomarker panel.

### Cross-Validated XGBoost Rank Aggregation

XGBoost models were fitted across stratified cross-validation folds, using class-balanced sample weights. The number of folds was five in most datasets and capped at the minimum class size where necessary. Within each fold, probes were ranked according to gain-based feature importance. The fold-specific rankings were aggregated using the mean rank and mean importance of each probe, prioritizing features that were consistently informative across alternative data partitions. The 500 highest-ranked probes were retained for subsequent RFECV refinement. A common XGBoost configuration was applied across all datasets because the model was used for feature screening rather than final prediction (Chen and Guestrin, 2016). Alternative candidate-set sizes were examined in the sensitivity analysis.

### SVM-RFECV Refinement

The candidate probes were refined using RFECV with a class-weighted linear SVM estimator. Balanced accuracy was used as the selection criterion, and stratified cross-validation was adapted by capping the number of folds to the size of the smallest class. At each iteration, 10% of the remaining probes were removed until the subset producing the highest mean cross-validation score was identified. Mean imputation and feature standardization were estimated within the corresponding cross-validation procedure. Feature-selection consistency was summarized using fold-level selection frequencies, and probes retained in at least 50% of folds were designated as consensus features.

### Post-Selection Panel Assessment

The final feature panel identified for each dataset was assessed using five repeats of stratified cross-validation, with the number of folds reduced when necessary according to the size of the smallest class. Logistic regression was selected as the primary model because it provides a simple linear benchmark for the selected gene panels and avoids presenting biomarker discovery as dependent on a highly complex classifier. As a secondary comparator, we used a stacking ensemble comprising logistic regression, support vector machine, random forest, and XGBoost as base classifiers and logistic regression as the meta-classifier. The meta-classifier was trained using cross-validated class-probability predictions generated for all training samples, with each prediction obtained from a base classifier that had not been trained on the corresponding sample. Results for the individual classifiers and soft-voting ensemble were retained as supplementary analyses.

For each repeated cross-validation split, model preprocessing was fitted only on the training portion of that split. Base learners in the stacking classifier were wrapped in imputation and scaling pipelines where appropriate. The stacking meta-learner was trained using out-of-fold predictions generated within the training portion of each repeated cross-validation split, as implemented by scikit-learn’s `StackingClassifier`. Additional classifiers, including SVM, random forest, XGBoost, and soft voting, were retained as supplementary robustness analyses.

Performance was evaluated using accuracy, balanced accuracy, macro and weighted F1 scores, macro precision, macro recall, Matthews correlation coefficient, ROC–AUC, and average precision where applicable. Weighted F1, balanced accuracy, and Matthews correlation coefficient were emphasized in the main text because they are more informative than accuracy alone for datasets with unequal class distributions.

Because feature selection was performed using each complete dataset before this evaluation, the reported scores represent the internal discriminative capacity of the selected panels. They should not be interpreted as unbiased estimates of complete-pipeline performance on independent samples or as external clinical validation. The absence of external test cohorts is addressed explicitly as a limitation.

### SHAP-Based Interpretation

SHapley Additive exPlanations (SHAP) were used to examine how the selected features contributed to model predictions (Lundberg and Lee, 2017). For each dataset, an XGBoost interpretation model was fitted using the final selected or consensus feature panel, and mean absolute SHAP values were calculated to summarize global feature importance. For binary datasets, features were ranked according to their overall contribution to class discrimination; for multiclass datasets, class-specific SHAP summaries were generated to identify features associated with individual subtypes. SHAP values were interpreted as measures of predictive contribution rather than evidence of causal biological effects. Probe identifiers were mapped to gene symbols where annotations were available; unmapped probes retained their original identifiers, and multiple probes mapping to the same gene were kept as separate features unless collapsed for visualization.

### Enrichment Analysis

Functional enrichment analysis was performed on selected consensus gene sets for representative datasets with sufficiently large and biologically interpretable gene lists. Probe-level features were converted to available gene symbols before enrichment analysis; unmapped probes were retained in feature tables and SHAP outputs but were not used as gene symbols for enrichment queries. When multiple probes mapped to the same gene symbol, the gene symbol was represented once in the submitted enrichment list to avoid inflating term counts.

Enrichment was explored using Enrichr (Chen et al., 2013; Kuleshov et al., 2016; Xie et al., 2021), g:Profiler (Kolberg et al., 2023), DAVID (Sherman et al., 2022; Huang et al., 2009), Metascape (Zhou et al., 2019), and ToppGene (Chen et al., 2007). For each tool, the default background universe was used, and enriched terms were summarized using Benjamini-Hochberg adjusted significance values where available. Because background definitions, annotation coverage, and implementation details differ across enrichment platforms, the resulting terms were interpreted as convergent biological context rather than as a formal cross-tool statistical meta-analysis. Enrichment analysis was not applied to very small signatures, such as the five-gene leukemia panel, because pathway-level tests are underpowered and difficult to interpret for very short gene lists. Enrichment results were used as supportive biological context rather than as proof of causal disease mechanisms.

### Implementation

All analyses were implemented in Python using scikit-learn, XGBoost, pandas, NumPy, SHAP-compatible XGBoost contribution outputs, and plotting libraries. The base random seed was fixed at 42. Stratified XGBoost and RFECV splitters used shuffling with `random state=42`; fold-specific XGBoost models used seeds 42 plus the one-based fold index. Logistic regression, SVM, random forest, XGBoost, and the stacking meta-learner were also constructed with fixed random states. Aggregated XGBoost features were ordered deterministically by ascending mean rank, descending mean importance, and feature identifier. These settings make the workflow reproducible for identical ordered input matrices and the same software environment. Per-task configuration files record the seed, requested and adaptive fold counts, selected-feature counts, and final parameter settings.

The workflow produced XGBoost fold-wise feature rankings, aggregated feature ranks, RFECV-selected panels, fold-level RFECV selections, consensus feature tables, repeated cross-validation metrics, SHAP summaries, and enrichment-ready gene lists for downstream analysis (refer to Figure 1.

**Figure 1.**
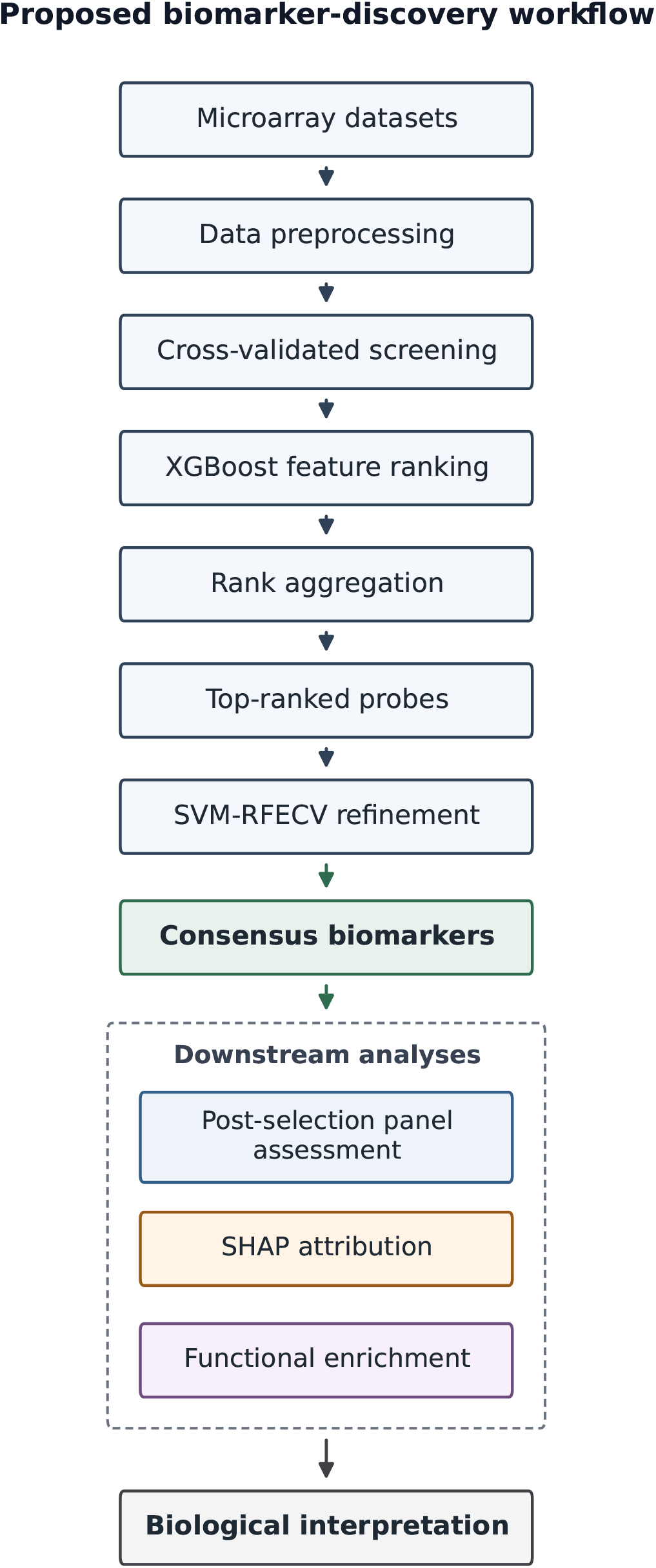
Overview of the proposed biomarker-discovery workflow. Cross-validated XGBoost ranking reduces the feature space, and SVM-RFECV refines the top-ranked probes into consensus candidate biomarkers. The resulting panels undergo post-selection assessment, model-specific SHAP attribution, and functional enrichment to support biological interpretation. Performance assessment occurs after full-cohort feature selection.

### Parameter Settings and Sensitivity Analysis

Final parameter choices were fixed before producing the manuscript outputs. XGBoost rank aggregation retained the top 500 probes for RFECV refinement, using fixed screening hyperparameters across all datasets. RFECV used a linear SVM estimator, balanced accuracy scoring, class-balanced weights, and an elimination step of 0.10. The nominal fold count was five for XGBoost screening, RFECV, and repeated panel assessment. It was reduced to three for endometrial cancer and four for Leukemia (4-class), matching the corresponding minimum class sizes. Repeated panel assessment used five repeats, and all manuscript outputs used the base seed 42. The final configuration used for the manuscript outputs is summarized in Table 1.

**Table 1.** Final configuration used for manuscript outputs.

| Component | Final setting | Sensitivity range or note |
| --- | --- | --- |
| XGBoost screening | Top 500 ranked probes | Top 200, 300, 500, and 5%–20% candidate cutoffs explored |
| XGBoost hyperparameters | 200 trees, depth 3, learning rate 0.05, subsample 0.8, column subsample 0.8, gain importance | Fixed across datasets; no dataset-specific tuning |
| RFECV estimator | Linear SVM | Chosen for high-dimensional data and interpretable feature weights |
| RFECV scoring | Balanced accuracy | Used to reduce dominance of majority classes |
| RFECV elimination step | 0.10 | Steps from 0.05 to 0.50 explored |
| Cross-validation folds | Stratified and adapted to smallest class size | Used when minority class sizes were smaller than the nominal fold count |
| Randomness control | Base seed 42 for splitters and stochastic estimators | Fold-specific XGBoost seeds derived deterministically from the base seed |
| Panel assessment | Five repeats of stratified cross-validation | Applied after full-cohort selection to the fixed final panels |
| Consensus criterion | Selected in at least 50% of RFECV folds | Used for SHAP and enrichment analysis when biologically suitable |

Preliminary sensitivity analyses were used to compare candidate screening sizes and RFECV elimination steps. XGBoost candidate sets were explored using fixed top-ranked probe counts of 200, 300, and 500, as well as percentage-based cutoffs from 5% to 20% of ranked probes. RFECV elimination steps from 0.05 to 0.50 were also examined. The final settings were selected as a practical compromise between predictive performance, panel compactness, and runtime. These sensitivity analyses were treated as configuration checks rather than a separate benchmark because the study goal was to derive interpretable candidate biomarker panels from small public datasets.

## Results

### Dimensionality Reduction and Selected Feature Panels

The two-stage feature-selection workflow reduced the original dimensionality of all 21 datasets. The initial expression matrices contained between 1,626 and 54,613 probe-level features, whereas the final SVM-RFECV panels contained between 5 and 300 features. Among the binary datasets, panel sizes ranged from 5 to 250 features, with the smallest panel obtained for the leukemia ALL-versus-AML dataset. Among the multiclass datasets, panel sizes ranged from 50 to 300 features, with the largest panel selected for the multiclass lung cancer dataset. Dataset-specific sample sizes, original feature counts, and final panel sizes are reported in Table 2.

**Table 2.** Post-selection repeated cross-validation performance summary for all 21 classification tasks.

| Dataset | Task | Samples | Classes | Original features | Selected features | LR weighted $F_1$ | LR balanced accuracy | LR MCC | Stacking weighted $F_1$ | Stacking balanced accuracy | Stacking MCC |
| --- | --- | --- | --- | --- | --- | --- | --- | --- | --- | --- | --- |
| Adenocarcinoma | Binary | 36 | Tumor(18), Normal(18) | 7457 | 50 | 1.00 | 1.00 | 1.00 | 1.00 | 1.00 | 1.00 |
| Brain tumor | Binary | 60 | Tumor(21), Normal(39) | 7129 | 150 | 0.92 | 0.93 | 0.86 | 0.91 | 0.92 | 0.83 |
| Breast cancer | Binary | 97 | Relapse(46), Non-relapse(51) | 24481 | 100 | 0.96 | 0.96 | 0.92 | 0.95 | 0.95 | 0.90 |
| Colon tumor | Binary | 62 | Tumor(40), Normal(22) | 2000 | 200 | 0.90 | 0.90 | 0.80 | 0.90 | 0.89 | 0.79 |
| Gastric cancer | Binary | 65 | Tumors(29), non-malignants(36) | 22645 | 100 | 1.00 | 1.00 | 1.00 | 0.99 | 0.99 | 0.98 |
| Leukemia | Binary | 72 | ALL(47), AML(25) | 7129 | 5 | 0.98 | 0.99 | 0.97 | 0.97 | 0.96 | 0.93 |
| Lung cancer | Binary | 181 | MPM(31), AD(150) | 1626 | 150 | 1.00 | 1.00 | 1.00 | 1.00 | 1.00 | 1.00 |
| Lymphoma | Binary | 77 | DLBCL(58), FL(19) | 2647 | 100 | 0.97 | 0.98 | 0.93 | 1.00 | 1.00 | 0.99 |
| Myeloma | Binary | 173 | Presence(137) and absence(36) of focal lesions of bone | 12625 | 100 | 1.00 | 1.00 | 0.99 | 0.99 | 0.99 | 0.98 |
| Ovarian cancer | Binary | 253 | Cancer(162), Normal(91) | 15154 | 50 | 1.00 | 1.00 | 1.00 | 1.00 | 1.00 | 1.00 |
| Prostate cancer | Binary | 102 | Tumor(52), Normal(50) | 2135 | 250 | 0.97 | 0.97 | 0.94 | 0.95 | 0.95 | 0.90 |
| Brain cancer | Multiclass | 130 | Ependymoma(46), glioblastoma(34), medulloblastoma(22), normal(13), pilocytic astrocytoma(15) | 16382 | 100 | 0.98 | 0.98 | 0.97 | 0.97 | 0.97 | 0.97 |
| Crohn's disease | Multiclass | 126 | 1(59), 2(41), 3(26) | 22283 | 100 | 0.99 | 0.99 | 0.98 | 0.99 | 0.99 | 0.99 |
| Endometrial cancer | Multiclass | 42 | PS(13), CC(3), E(19), N(7) | 1771 | 200 | 0.85 | 0.77 | 0.81 | 0.83 | 0.75 | 0.78 |
| Glioma | Multiclass | 180 | 1(26), 2(81), 3(23), 4(50) | 54613 | 50 | 0.88 | 0.88 | 0.82 | 0.88 | 0.89 | 0.83 |
| Leukemia (3-class) | Multiclass | 72 | B-cell(38), T-cell(9), AML(25) | 7129 | 50 | 0.99 | 0.99 | 0.98 | 0.98 | 0.97 | 0.97 |
| Leukemia (4-class) | Multiclass | 72 | B-cell(38), BM(21), PB(4), T-cell(9) | 7129 | 50 | 0.97 | 0.93 | 0.96 | 0.96 | 0.93 | 0.95 |
| Lung cancer | Multiclass | 203 | 1(139), 2(17), 3(6), 4(21), 5(20) | 12600 | 300 | 0.97 | 0.97 | 0.95 | 0.98 | 0.97 | 0.96 |
| Lymphoma | Multiclass | 66 | DLBCL(46), FL(9), CLL(11) | 4026 | 50 | 1.00 | 1.00 | 1.00 | 1.00 | 1.00 | 1.00 |
| MLL | Multiclass | 72 | ALL(24), MLL(20), AML(28) | 12582 | 100 | 1.00 | 1.00 | 1.00 | 0.99 | 0.99 | 0.99 |
| SRBCT | Multiclass | 83 | 1(29), 2(11), 3(18), 4(25) | 2308 | 50 | 1.00 | 1.00 | 1.00 | 1.00 | 1.00 | 1.00 |

Fold-level feature selections were also summarized to examine consistency across RFECV partitions. For each dataset, the selection frequency of every retained feature was calculated, and features appearing in at least 50% of the RFECV folds were designated as consensus features. These consensus sets were used for subsequent SHAP interpretation and enrichment analysis where sufficient annotated features were available. Complete selected panels and fold-level selection frequencies are provided in the Supplementary Material.

### Post-Selection Classification Assessment

The selected gene panels were evaluated using repeated stratified cross-validation. Logistic regression was used as the primary reported classifier because it provides a parsimonious linear assessment of the selected panels, while stacking was included as a secondary ensemble comparator. The main performance results are summarized in Table 2, and the fold-level weighted F_1_ distributions are shown in Figure 2.

**Figure 2.**
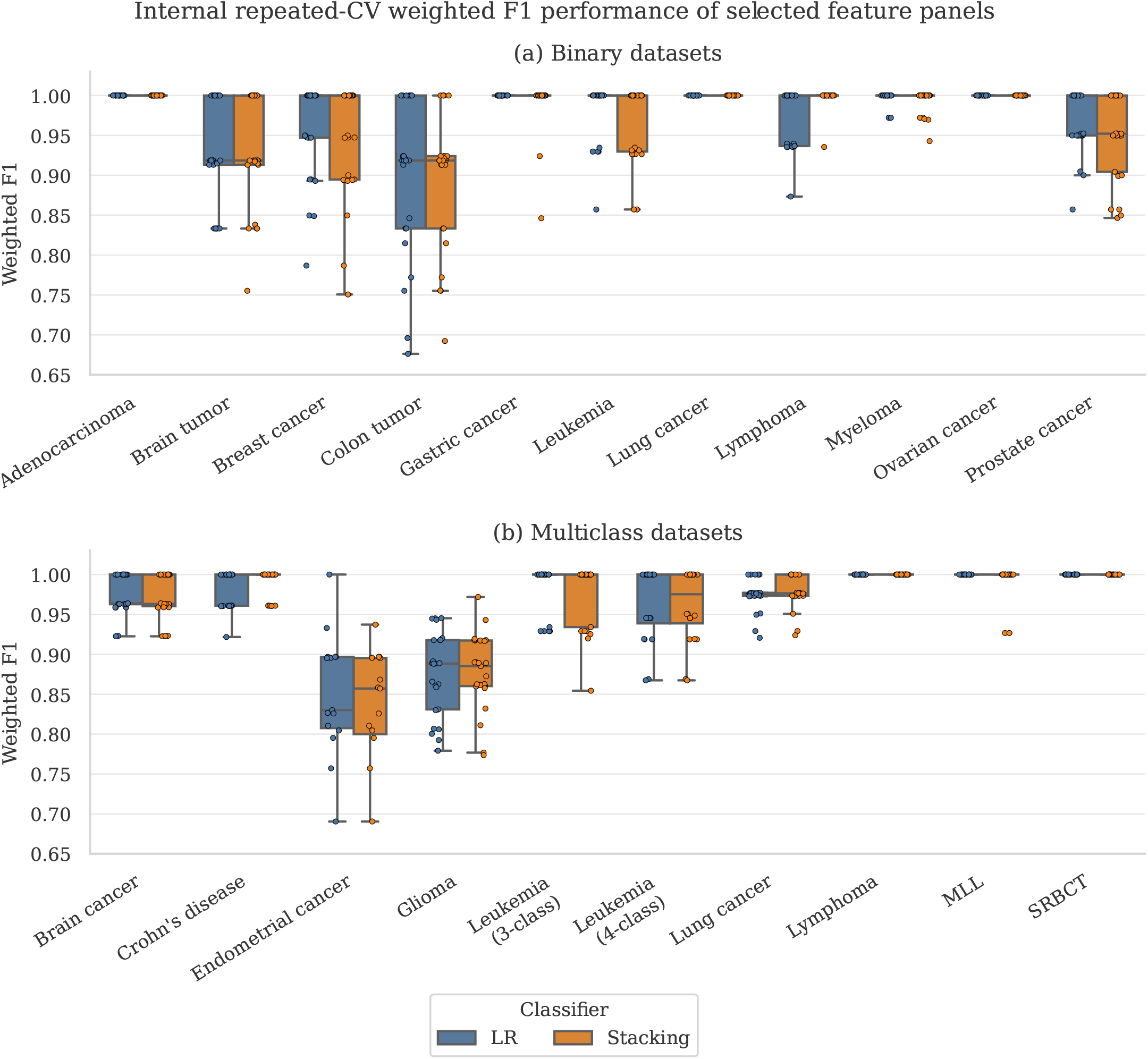
Post-selection weighted F1 distribution across classification tasks.** Boxplots summarize repeated cross-validation weighted F1 scores for logistic regression and stacking on the fixed selected panels, with binary and multiclass tasks shown as separate panels.

Across the 11 binary datasets, logistic regression achieved weighted F1 scores ranging from 0.90 to 1.00. Perfect weighted F1 and balanced accuracy were obtained for adenocarcinoma, gastric cancer, lung cancer, and ovarian cancer, while myeloma also achieved near-perfect performance. Colon tumor produced the lowest binary weighted F1 score at 0.90, followed by brain tumor at 0.92. The five-feature leukemia panel achieved a weighted F1 score of 0.98, balanced accuracy of 0.99, and MCC of 0.97.

For the 10 multiclass datasets, logistic-regression weighted F1 scores ranged from 0.85 to 1.00. Multiclass lymphoma, MLL, and SRBCT achieved perfect weighted F1 scores, whereas endometrial cancer and glioma produced the lowest values, at 0.85 and 0.88, respectively. The stacking ensemble generally performed similarly to logistic regression, providing modest improvements for some datasets, including binary lymphoma and multiclass lung cancer, but not consistently outperforming the linear model. Overall, the limited improvement provided by stacking indicates that the selected panels retained substantial discriminative information that could be captured using a comparatively simple classifier.

### Contextual Comparison with Previous Microarray Studies

Figure 3 places the internal classification accuracies obtained in this study alongside values reported for overlapping datasets in previous microarray feature-selection studies. Across both binary and multiclass tasks, logistic regression and stacking generally produced results within the range of published methods, with several datasets lying near the upper end of the reported values.

**Figure 3.**
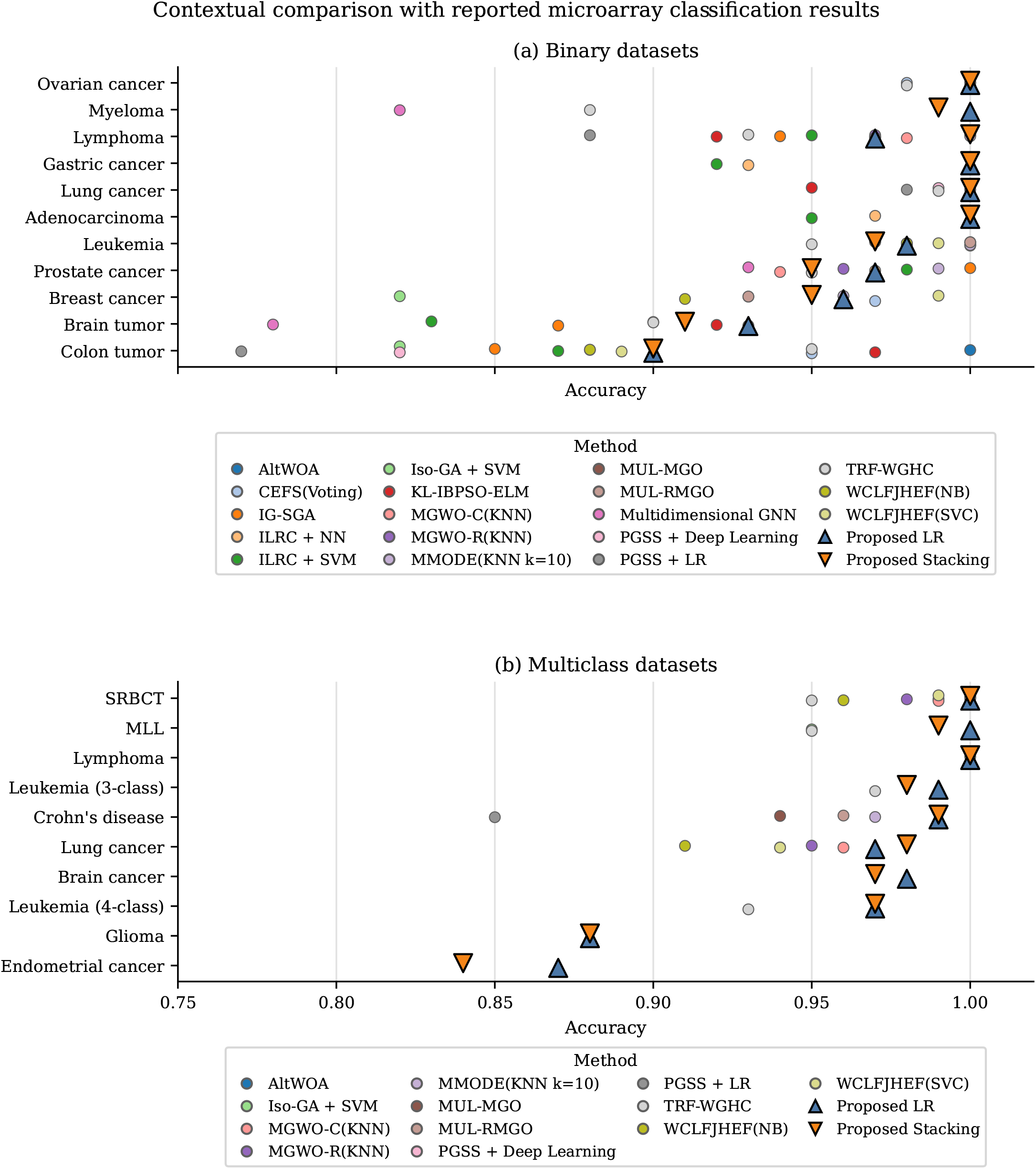
Contextual comparison of the proposed method with reported microarray classification results. Colored circular markers denote accuracies reported by individual methods in literature, with method-specific legends shown for binary and multiclass panels. Triangle markers denote the proposed logistic regression and stacking results. The diagram is presented as literature context rather than as a fully controlled benchmark because the reported values come from studies using heterogeneous validation protocols, preprocessing strategies, feature-selection settings, and classifiers.

These comparisons should not be interpreted as a controlled benchmark or evidence of statistical superiority. The published studies differed in preprocessing, selected feature counts, classifiers, cross-validation designs, and performance-reporting procedures. Cross-validation was also the predominant evaluation strategy among the representative studies, reflecting the limited sample sizes of commonly used microarray datasets; nevertheless, differences in fold structure and in whether feature selection was repeated within each validation fold limit direct comparison of the reported accuracies. Accuracy was therefore used only because it was the metric most consistently available across the reviewed studies. Figure 3 is presented to provide literature context rather than a direct head-to-head evaluation.

### SHAP-based Interpretation of Selected Signatures

SHAP analysis was used to examine the contribution of selected genes to model discrimination in representative datasets. Because SHAP values quantify feature contribution to model output, these analyses were interpreted as evidence of class-discriminative relevance rather than causal biological effects. For binary datasets, mean absolute SHAP values were used to rank the selected genes by their overall contribution to class separation. For multiclass datasets, classwise SHAP summaries were used to identify features contributing preferentially to individual classes.

In the breast cancer relapse dataset, the highest SHAP contributions were observed for *TSPYL5, SCUBE2, FGD6*, and *H4F5* (Figure 4a). These features dominated the relapse/non-relapse attribution profile, indicating that the classifier relied on a compact set of discriminative genes rather than a diffuse contribution across the full selected panel. TSPYL5 has been associated with poor breast cancer outcomes and metastatic progression through regulation of p53- and PTEN-related pathways (Epping et al., 2011; Shi et al., 2026), and SCUBE2 has been implicated in bone metastasis in luminal breast cancer (Wu et al., 2023), whereas the relevance of FGD6 and H4F5 remains less established. The signature therefore provides biologically plausible, but hypothesis-generating, candidates for further validation.

**Figure 4.**
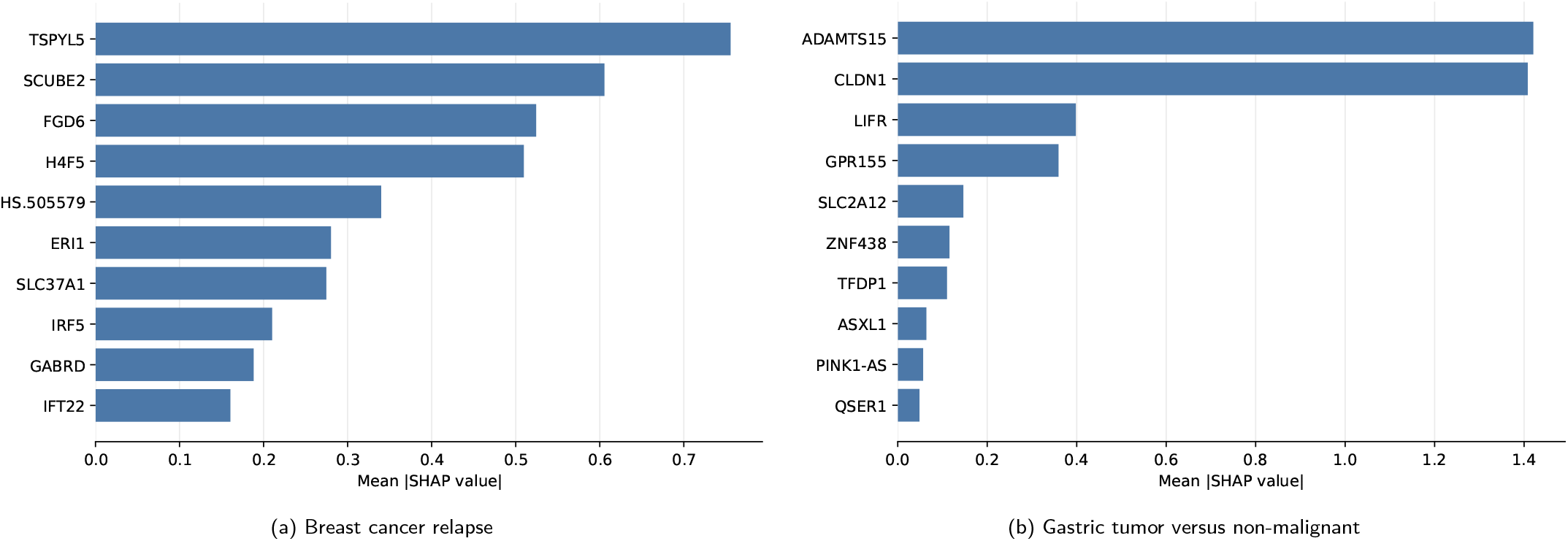
Representative binary SHAP summaries. Bars indicate mean absolute SHAP values for the highest-contributing selected genes. These values quantify contribution magnitude and do not by themselves indicate the direction of class association.

In the gastric tumor versus non-malignant dataset, the attribution profile was dominated by *ADAMTS15* and *CLDN1*, with additional contributions from *LIFR* and *GPR155* (Figure 4b). CLDN1 is frequently upregulated in gastric adenocarcinoma and has been associated with reduced postoperative survival (Eftang et al., 2013), while LIFR has been implicated in gastric cancer progression and poor prognosis (Park et al., 2025). Reduced GPR155 expression has been associated with hematogenous metastasis and recurrence (Shimizu et al., 2017). In one study, ADAMTS15 has been included in a serum protein signature with high diagnostic performance for resectable gastric cancer, providing supportive, although not gene-specific, evidence for its relevance (Shen et al., 2019). Together, these features suggest that the predictive signature may reflect processes involving extracellular-matrix regulation, epithelial organization, cytokine signaling, and metastatic potential. However, their SHAP contributions indicate predictive relevance rather than causal or coordinated biological activity, and further experimental validation is required.

For the multiclass brain cancer dataset, classwise SHAP analysis revealed subtype-specific feature contributions (Figure 5). *PHF21B* contributed predominantly to medulloblastoma classification, while *CASKIN1* contributed mainly to the normal class. Ependymoma classification was influenced by OPG-associated probes and *MSX1*, glioblastoma was influenced strongly by *GDF10*, and pilocytic astrocytoma showed contributions from *GGH* and additional probe-level features. These findings indicate that the multiclass classifier used class-specific molecular signals rather than a single shared cancer signature.

**Figure 5.**
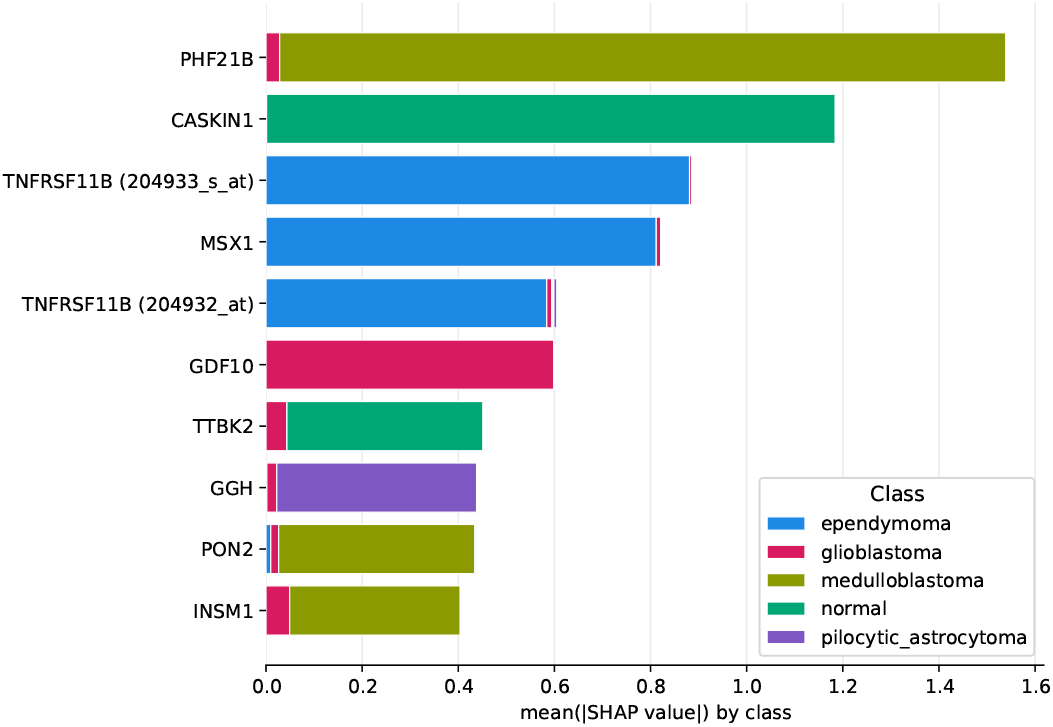
Classwise SHAP summary for the multiclass brain cancer dataset. Bars represent mean absolute SHAP contribution by class, highlighting subtype-specific features used by the model.

**Figure 6.**
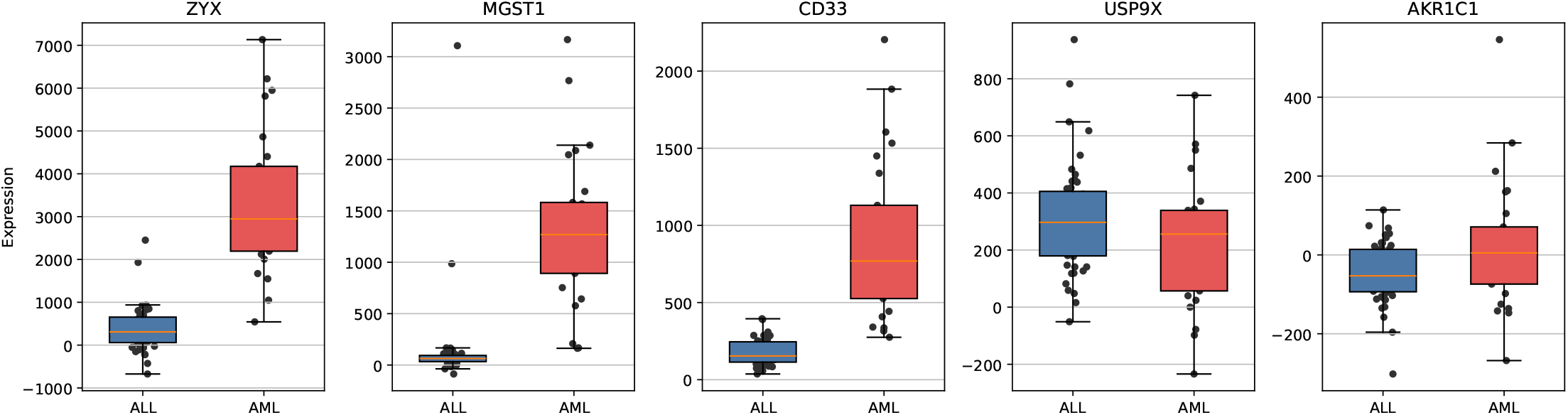
Classwise expression patterns of the five-gene leukemia signature. The selected genes show varying degrees of separation between ALL and AML samples, with *CD33, MGST1*, and *ZYX* providing particularly clear class-associated patterns.

**Figure 7.**
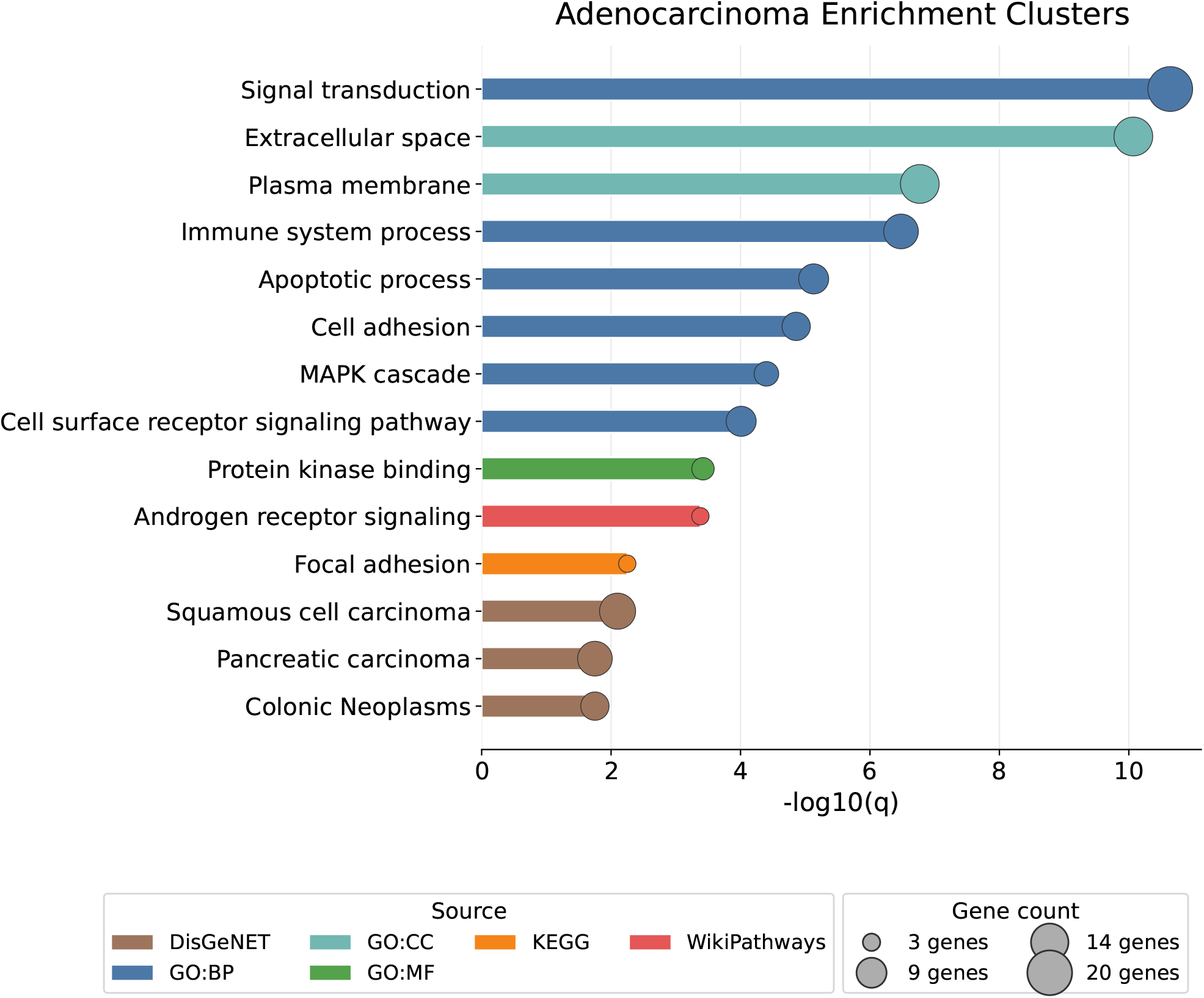
Functional enrichment analysis of the adenocarcinoma consensus signature. Bars indicate enrichment significance as *−* log_10_(*q*), while point size represents the number of genes contributing to each term. Enrichment results are interpreted as supportive biological context for the selected signature.

### Biological relevance of selected gene signatures

To assess whether the selected signatures reflected biologically meaningful disease signals, we examined representative gene-level and pathway-level evidence. Because selected gene panels varied considerably in size, biological relevance was assessed using different strategies depending on the dataset. For very small signatures, gene-level interpretation was emphasized; for larger consensus signatures, pathway enrichment analysis was used to provide broader functional context.

The five-feature signature distinguished AML from ALL, consistent with foundational studies demonstrating that acute leukemia classes can be separated using gene-expression profiles (Golub et al., 1999; Thomas et al., 2001). The inclusion of CD33 provides strong biological plausibility for AML–ALL discrimination. CD33 is an established myeloid differentiation antigen and therapeutic target in AML, and circulating CD33 concentrations have been reported to be significantly higher in AML than in ALL and healthy controls (Abdool et al., 2010; Walter et al., 2012). Nevertheless, its contribution should be interpreted within the complete five-feature signature rather than as evidence that CD33 alone is sufficient for leukemia classification. USP9X has been shown to promote AML-cell survival by stabilizing the RNA *m*^6^*A* demethylase ALKBH5 (Wang et al., 2023). These associations provide biological context for the selected panel but do not establish that its members jointly regulate lymphoid–myeloid lineage differences.

For adenocarcinoma, the selected consensus gene set was large enough to support pathway-level interpretation. Functional enrichment of the 36-gene colorectal adenocarcinoma signature highlighted processes related to signal transduction and MAPK signalling, extracellular and plasma-membrane organisation, cell adhesion and focal adhesion, immune responses, and apoptosis. These categories are consistent with established biological processes involved in colorectal-cancer development, although enrichment alone does not demonstrate pathway activation. Individual signature members provided further biological context. SOX9 and KRT20 have been used as markers of aberrant stem-like activity and intestinal differentiation, respectively, in colorectal-cancer models, suggesting that the signature captures variation in epithelial cell state (Spisak et al., 2024). MMP7 has been linked experimentally to colorectal-cancer-cell migration and adverse clinical features (Ou et al., 2023), whereas LTBP2 has been identified in an extracellular-matrix-remodelling cancer-associated fibroblast population in colorectal liver metastases (Giguelay et. al., 2022). HMGB1 has also been associated with colorectal neoplastic progression and an immune-cold tumour phenotype (Porter et al., 2023). Collectively, these findings suggest that the signature reflects both tumour-epithelial and microenvironment-related processes. Nevertheless, the identified associations provide biological support rather than evidence of causal mechanisms or formal consensus molecular subtype classification. The enrichment of the signature for colonic neoplasms further supports its disease relevance, although disease-term enrichment should not be interpreted as independent diagnostic validation.

Overall, the biological relevance analyses suggest that the proposed pipeline can identify signatures with both discriminative and interpretable properties. Gene-level interpretation was most appropriate for very compact signatures such as leukemia, while enrichment analysis provided pathway-level support for larger consensus gene sets such as adenocarcinoma. These analyses were treated as supportive biological context and not as independent validation of causal disease mechanisms.

### Parameter Sensitivity and Feature Stability

Preliminary parameter exploration indicated that overly broad XGBoost candidate sets tended to increase the size of the final RFECV panels without consistently improving repeated cross-validation performance. Conversely, very small candidate sets sometimes removed useful complementary probes before wrapper refinement. Across the explored settings, which included top-200, top-300, top-500, and 5%-20% XGBoost candidate cutoffs, retaining the top 500 XGBoost-ranked probes provided a stable working point across the dataset collection. Similarly, an RFECV elimination step of 0.10 was selected after examining step sizes from 0.05 to 0.50 because it reduced runtime without producing excessively coarse feature-removal behavior.

Feature-selection stability was summarized using RFECV fold-level selection frequencies. Genes selected in at least 50% of RFECV folds were treated as consensus features for SHAP interpretation and enrichment analysis where appropriate. This consensus criterion was used to reduce dependence on a single RFECV split and to prioritize recurrently selected genes.

As an additional stability check, pairwise Jaccard similarity was calculated among the RFECV fold-selected feature sets for all 21 datasets. Across the 21 datasets, mean pairwise Jaccard similarity ranged from 0.218 to 0.720, with the highest fold-level agreement observed for multiclass lung cancer and the lowest for glioma. These results indicate that selection stability varied substantially among datasets: several signatures showed moderate fold-to-fold overlap, whereas others were more sensitive to the composition of the training samples. Given the limited sample sizes and small minority classes in several datasets, the stability estimates were treated as supplementary evidence rather than definitive confirmation of biomarker reproducibility.

### Summary of Main Findings

Overall, the results suggest that the proposed two-stage pipeline can identify compact gene panels with strong internal discriminative capacity across diverse binary and multiclass microarray datasets. Logistic regression performed strongly on the selected panels, indicating that many signatures retained linearly separable class information. SHAP analysis provided gene-level interpretability, while enrichment analysis supplied pathway-level biological context for representative consensus signatures. These findings support the use of the pipeline as a practical biomarker-discovery framework for small-sample microarray studies, with the caveat that external validation remains necessary before clinical translation.

## Discussion

This study evaluated an interpretable two-stage workflow for biomarker discovery from small-sample, high-dimensional microarray datasets. Cross-validated XGBoost ranking provided an efficient initial reduction of the probe space, while SVM-RFECV refined the retained candidates into dataset-specific feature panels. Across 21 binary and multiclass datasets, the resulting panels preserved substantial class-discriminative information under repeated internal validation. Their strong performance with logistic regression further indicates that the retained information was not dependent exclusively on complex nonlinear classifiers. Together with SHAP attribution, consensus selection summaries, and functional enrichment, these results support the workflow as a practical framework for generating interpretable candidate biomarker signatures rather than as a new classification algorithm or a clinically validated diagnostic system.

The two-stage design addresses an important computational trade-off in microarray feature selection. Applying wrapper selection directly to tens of thousands of probes is computationally expensive and may increase sensitivity to sampling variation. The XGBoost stage therefore served as an embedded screening mechanism that reduced the search space before classifier-guided refinement. Similar filter–wrapper and embedded–wrapper combinations have been used in previous microarray studies because preliminary screening can make subsequent subset evaluation more tractable (Chandrashekar and Sahin, 2014; Got et al., 2021; Li et al., 2023). In the present workflow, fold-wise rank aggregation was used instead of relying on the importance values from a single fitted XGBoost model. Although this approach does not eliminate instability, it reduces dependence on one data partition and prioritizes probes that repeatedly obtain high rankings across folds.

An important observation was that logistic regression generally performed comparably to the stacking ensemble. This finding suggests that, after feature selection, many datasets contained class information that could be represented using relatively simple linear decision boundaries. In a biomarker-discovery context, this is advantageous because simpler predictive models provide a clearer assessment of whether a selected panel contains independently useful discriminative information. The stacking ensemble produced modest improvements for selected datasets but did not consistently outperform logistic regression, indicating that additional model complexity was not uniformly beneficial. Nevertheless, the high internal scores obtained for several small datasets should be interpreted cautiously, because near-perfect cross-validation performance can occur when class separation is strong, sample sizes are limited, or the selected panel has been derived using the complete cohort.

The biological analyses provided complementary levels of interpretation. SHAP attribution identified the selected features that contributed most strongly to model discrimination, whereas literature evidence supplied external biological context for several prioritized genes. In the breast cancer relapse dataset, the prominence of *TSPYL5* and *SCUBE2* was consistent with previous evidence linking these genes to aggressive breast cancer and metastatic behavior. Similarly, the gastric cancer signature contained genes with reported relevance to gastric malignancy, including *CLDN1, GPR155*, and *ADAMTS15*. The compact leukemia panel included *CD33*, an established myeloid antigen, and *USP9X*, which has been experimentally implicated in acute myeloid leukemia cell survival. These agreements support the biological plausibility of the selected signatures. However, SHAP values quantify contributions to a fitted prediction model and do not establish that the prioritized genes are causal disease drivers, independently diagnostic biomarkers, or members of a coordinated molecular mechanism.

Functional enrichment extended this interpretation from individual genes to broader biological processes. The adenocarcinoma consensus signature was enriched for signal transduction, extracellular and plasma-membrane processes, immune-system functions, apoptosis, cell adhesion, MAPK-related signaling, and focal adhesion. Several individual members of the signature also have reported associations with colorectal epithelial differentiation, invasion, extracellular-matrix organization, or the tumor microenvironment. The convergence of gene-level evidence and pathway-level enrichment suggests that the signature contains biologically coherent information related to both tumor cells and their surrounding microenvironment. Nevertheless, enrichment identifies statistical over-representation rather than pathway activation, and it does not demonstrate that the enriched processes directly caused the observed classification patterns. The use of default gene universes across enrichment tools also means that these results should be interpreted as supportive biological prioritization rather than definitive functional validation.

Feature-selection stability varied substantially across datasets. Pairwise Jaccard similarities ranged from 0.218 to 0.720, showing that some selected panels were relatively consistent across folds, whereas others were sensitive to changes in sample composition. This variability is expected in small-*n*, large-*p* data, where multiple correlated probes may provide similar predictive information and minor changes in the training samples can alter which member of a correlated group is selected. Consequently, a low Jaccard value does not necessarily imply an absence of biological signal, but it does indicate uncertainty regarding the exact composition of the proposed panel. Reporting fold-level selection frequencies and consensus signatures therefore provides more informative evidence than presenting a single feature list without an assessment of recurrence. The consensus threshold used in this study should nevertheless be regarded as a practical prioritization criterion rather than a formal guarantee of reproducibility.

The literature comparison showed that the internal accuracies obtained in this study were generally within the range reported by recent microarray feature-selection methods. However, these comparisons cannot support claims of statistical superiority because previous studies differed in preprocessing, feature-selection scope, classifier configuration, cross-validation design, and reported metrics. Some methods optimize classifiers and feature subsets jointly, whereas the present study used fixed screening settings and emphasized interpretable downstream analysis. The comparison should therefore be understood as contextual evidence that the workflow produces competitive internal discrimination while providing additional outputs—including consensus frequencies, SHAP attribution, and enrichment-ready gene lists—that support biological investigation. A controlled benchmark would require reimplementation of competing methods using identical data partitions, preprocessing procedures, metrics, and computational constraints.

Overall, the findings illustrate both the utility and the limitations of computational biomarker discovery from small public microarray cohorts. The workflow generated compact, interpretable panels across heterogeneous classification tasks, but the exact selected genes and the reported predictive scores remain dependent on the available samples, preprocessing history, platform annotations, and validation design. The panels should therefore be considered hypothesis-generating candidates for subsequent replication rather than final diagnostic or prognostic biomarkers. Independent cohort validation, preferably across laboratories or expression platforms, will be necessary to establish whether the prioritized signatures remain reproducible and biologically informative beyond the datasets from which they were derived.

### Limitations and Future Directions

Several considerations should be taken into account when interpreting the findings. First, the selected signatures were derived from relatively small public microarray cohorts and were not evaluated in independent external datasets. Repeated stratified cross-validation provided an internal assessment of the discriminative information retained by each panel, but validation across laboratories, patient populations, and expression platforms will be required before the signatures can be considered clinically generalizable.

Feature selection was performed using the complete dataset before repeated evaluation of the resulting panel. This design was chosen to derive a single interpretable signature from all available samples and to support downstream SHAP and enrichment analyses. Consequently, the reported performance values characterize the internal discriminative capacity of the selected panels and may be optimistic relative to performance on fully independent samples. Nested cross-validation would provide a stricter estimate of complete-pipeline generalization; however, in small-sample settings it may also produce high-variance estimates and heterogeneous feature panels across outer folds. Future work could therefore combine nested performance estimation with full-cohort refitting to obtain both an unbiased evaluation and a final consensus biomarker signature.

The study also relied on processed public expression matrices originating from different platforms and preprocessing procedures. Although the same downstream workflow was applied consistently, platform-specific normalization, probe annotation, and batch structure may influence the selected features. Cross-platform replication and updated probe-to-gene annotation would strengthen the biological interpretation of the panels.

Finally, SHAP attribution and functional enrichment provide interpretive evidence rather than causal validation. These analyses help prioritize biologically plausible candidates and pathways, but experimental studies are required to establish their mechanistic or clinical relevance.

## Conclusion

We developed an interpretable two-stage workflow for candidate biomarker discovery from small-sample microarray datasets.

Across 21 binary and multiclass tasks, XGBoost-based screening followed by SVM-RFECV produced substantially reduced feature panels that retained strong internal discriminative capacity. SHAP attribution, selection-frequency summaries, and functional enrichment provided complementary gene- and pathway-level context for representative signatures. The workflow offers a practical framework for generating biologically interpretable hypotheses from high-dimensional expression data; however, the selected panels require independent cohort replication and experimental validation before diagnostic, prognostic, or clinical interpretation.

## Supplementary Material

Supplementary tables include full dataset descriptions, complete selected feature lists, RFECV fold-level selected genes, RFECV Jaccard stability summaries, full repeated cross-validation metrics for all classifiers, model hyperparameters, and complete enrichment outputs. Supplementary figures include per-model performance plots, all SHAP summary plots, classwise SHAP plots for multiclass datasets, and additional expression plots for selected representative signatures.

## Data and Code Availability

The source code, processed outputs, plotting scripts, selected feature tables, SHAP outputs, and enrichment-ready gene lists are available at: https://github.com/mashiyat-mahjabin-prapty/microarray-feature-selection.

## Acknowledgements

Generative AI disclosure: OpenAI Codex was used to assist with language editing, code review, documentation, and consistency checking. All AI-assisted outputs were critically reviewed and verified by the authors, who take full responsibility for the manuscript, analysis, and conclusions.

## Author Contributions

Mashiyat Mahjabin Prapty performed the primary analysis, implemented the pipeline, interpreted the results, and drafted the manuscript. Mohammad Saifur Rahman supervised the study design, analysis, interpretation, and manuscript revision.

