## Supplementary material for "Interpretable biomarker discovery from small-sample microarray datasets using XGBoost rank aggregation and SVM-RFECV": Supplemenary_Materials

### Supplementary Methods

#### Preprocessing and adaptive validation

Expression matrices contained samples in rows and probe-level measurements in columns. Mean imputation was fitted inside applicable model-fitting procedures. Standardization was included inside the logistic-regression, SVM, and linear-SVM RFECV pipelines. Nominal five-fold stratification was capped at the smallest class count; repeated panel assessment used five repeats.

#### XGBoost rank aggregation and RFECV

Class-balanced XGBoost models ranked probes by gain across stratified folds. Fold rankings were aggregated deterministically by ascending mean rank, descending mean importance, and feature identifier. The final workflow retained the top 500 probes. RFECV used a class-weighted linear SVM, balanced accuracy, a 10% elimination step, and a minimum of five probes. Features retained in at least 50% of RFECV folds formed the consensus set.

#### Classifier configurations and stacking

The complete settings for LR, SVM, RF, XGBoost, voting, and stacking are listed in Table S2. For stacking, the base learners generated out-of-fold class-probability predictions within the training portion of each assessment split. A logistic-regression meta-classifier was fitted to those probabilities. The fitted stack was then evaluated on the untouched assessment fold.

#### Reproducibility

The base seed was 42. Shuffled splitters and stochastic estimators used fixed random states, and fold-specific XGBoost seeds were derived deterministically. The retained analysis environment records Python 3.12, NumPy 2.1.3, pandas 3.0.3, scikit-learn 1.9.0, XGBoost 3.3.0, SHAP 0.52.0, and Matplotlib 3.11.0.

#### Probe mapping and SHAP

Probe identifiers were mapped to gene symbols using the retained per-dataset mapping tables. Unmapped probes retained their probe identifiers. XGBoost prediction contributions were used for post-hoc SHAP interpretation. The retained numerical SHAP tables are gene-level aggregates; class-specific plots are supplied for multiclass tasks. Attribution values quantify predictive contribution and not causality.

#### Enrichment

Annotated representative consensus genes were examined using the enrichment tools described in the manuscript with their default background universes and adjusted significance values where available. Duplicate gene symbols were submitted once. The supplied enrichment export retained -log10(q) values, ontology source, overlap counts, and gene.

### Supplementary Tables

| **Table** | **Contents** |
| --- | --- |
| S1 Datasets | Dataset dimensions, class distributions, adaptive folds, selected-panel sizes, and repository-relative input locations. |
| S2 Parameters | Final workflow and model settings used for the manuscript outputs. |
| S3 Final Panels | Complete final feature panels and selection statistics. |
| S4 Stability | Pairwise Jaccard stability summaries across RFECV fold-specific selected sets. |
| S5 Performance Summary | Mean, standard deviation, and count for every classifier and metric. |
| S6 SHAP Rankings | Mean absolute SHAP rankings after probe-to-gene mapping. |
| S7 Enrichment | Representative adenocarcinoma functional-enrichment output. |
| S8 Literature | Related literature for comparison. |
| S9 Literature Comparison | Tidy data underlying the contextual literature comparison. |

### Supplementary Figures

| **Figure** | **Contents** |
| --- | --- |
| Figure S1 | RFECV fold-level feature-selection stability. |
| Figure S2 | SHAP bar and beeswarm summaries for all binary datasets. |
| Figure S3 | Overall and class-specific SHAP summaries for all multiclass datasets. |

**Figure S1**

RFECV fold-level stability

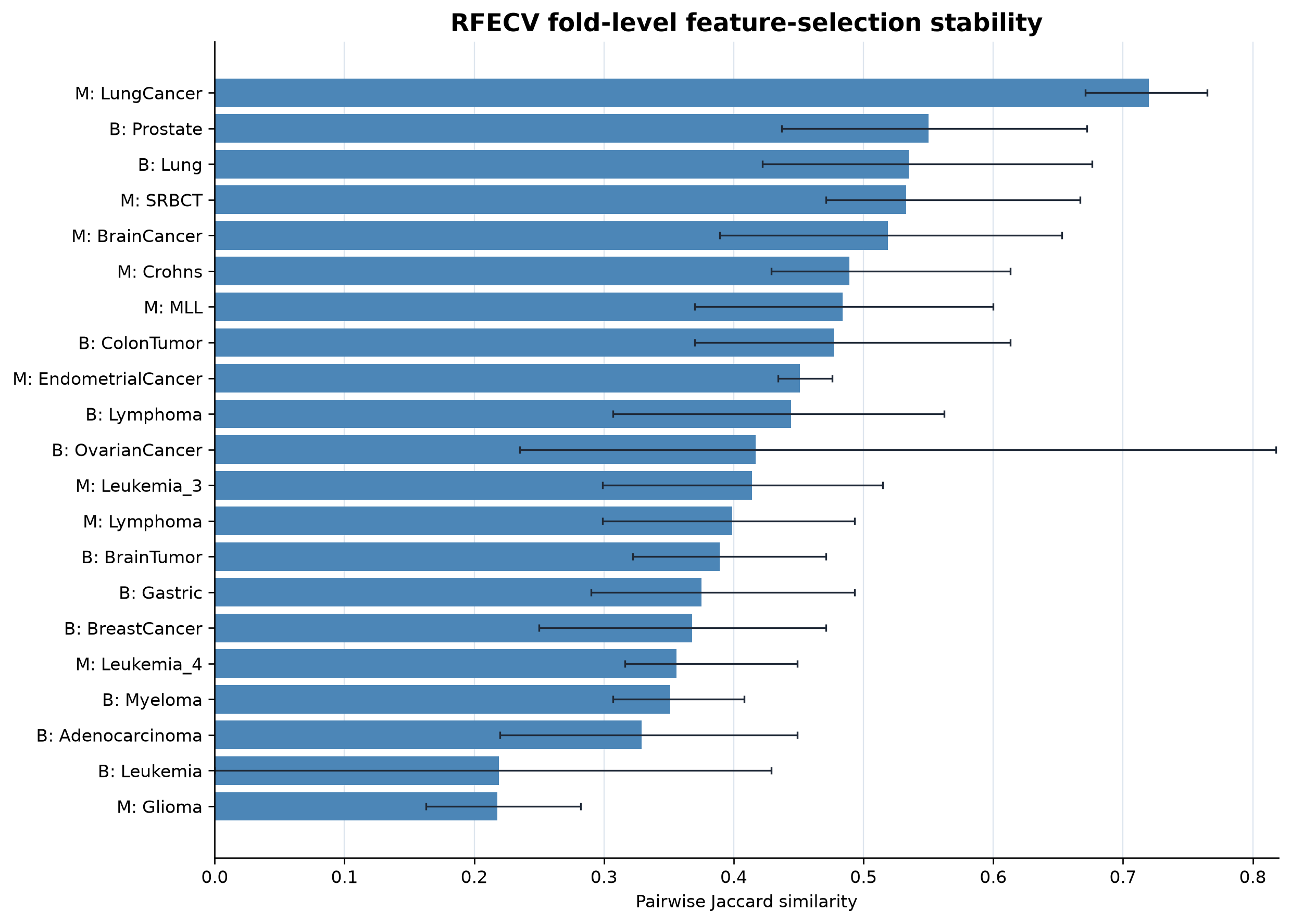

**Figure S1.** RFECV fold-level feature-selection stability. Bars show the mean pairwise Jaccard similarity for each dataset, and error bars span the observed minimum to maximum. B and M denote binary and multiclass tasks, respectively.

**Figure S2A**

Binary Adenocarcinoma

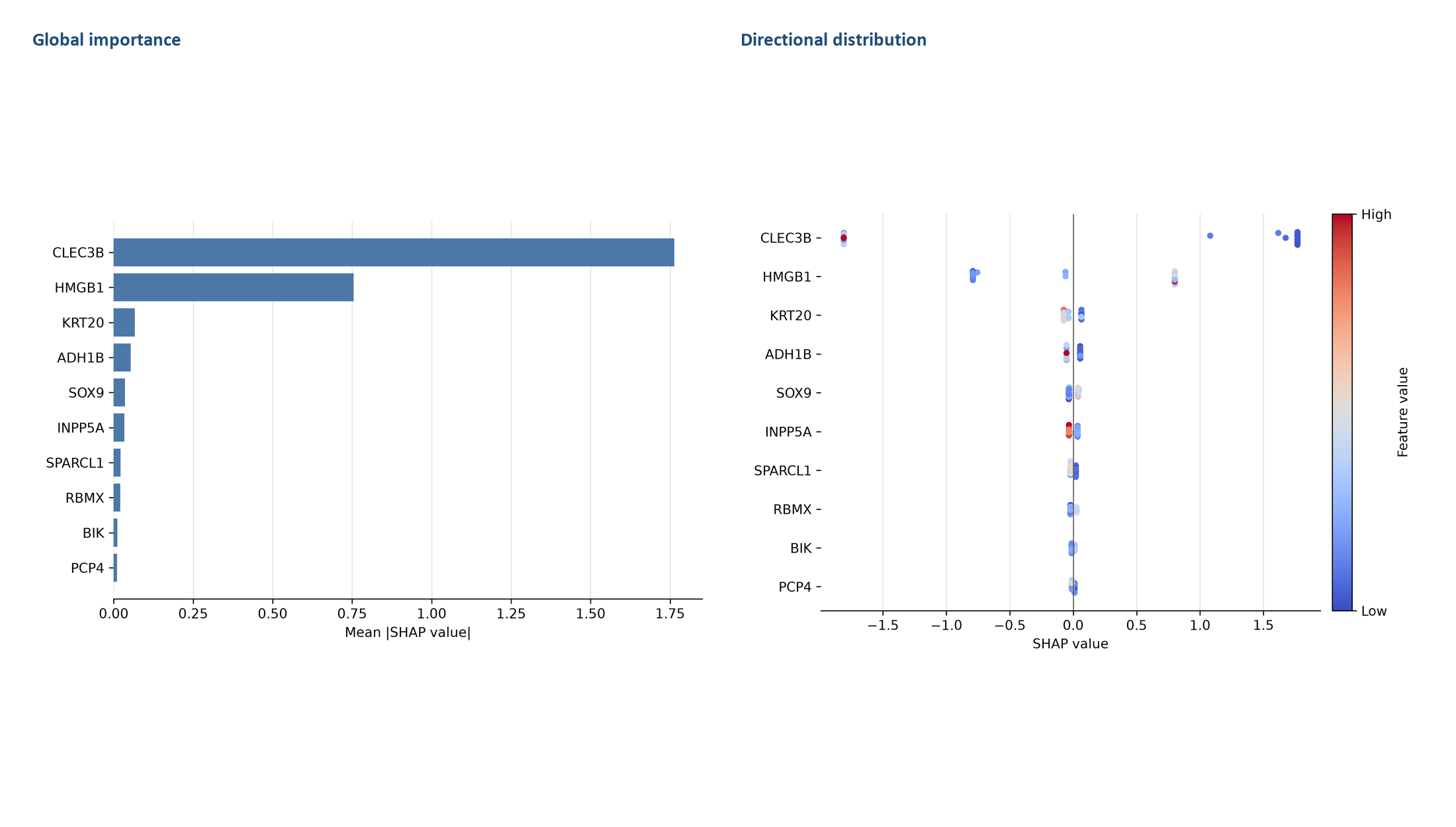

**Figure S2A.** SHAP summary for Binary Adenocarcinoma. The left panel ranks features by mean absolute SHAP value. The right panel shows the distribution and direction of feature contributions across samples; red indicates higher feature values and blue indicates lower values. SHAP values indicate predictive attribution, not causality.

**Figure S2B**

Binary BrainTumor

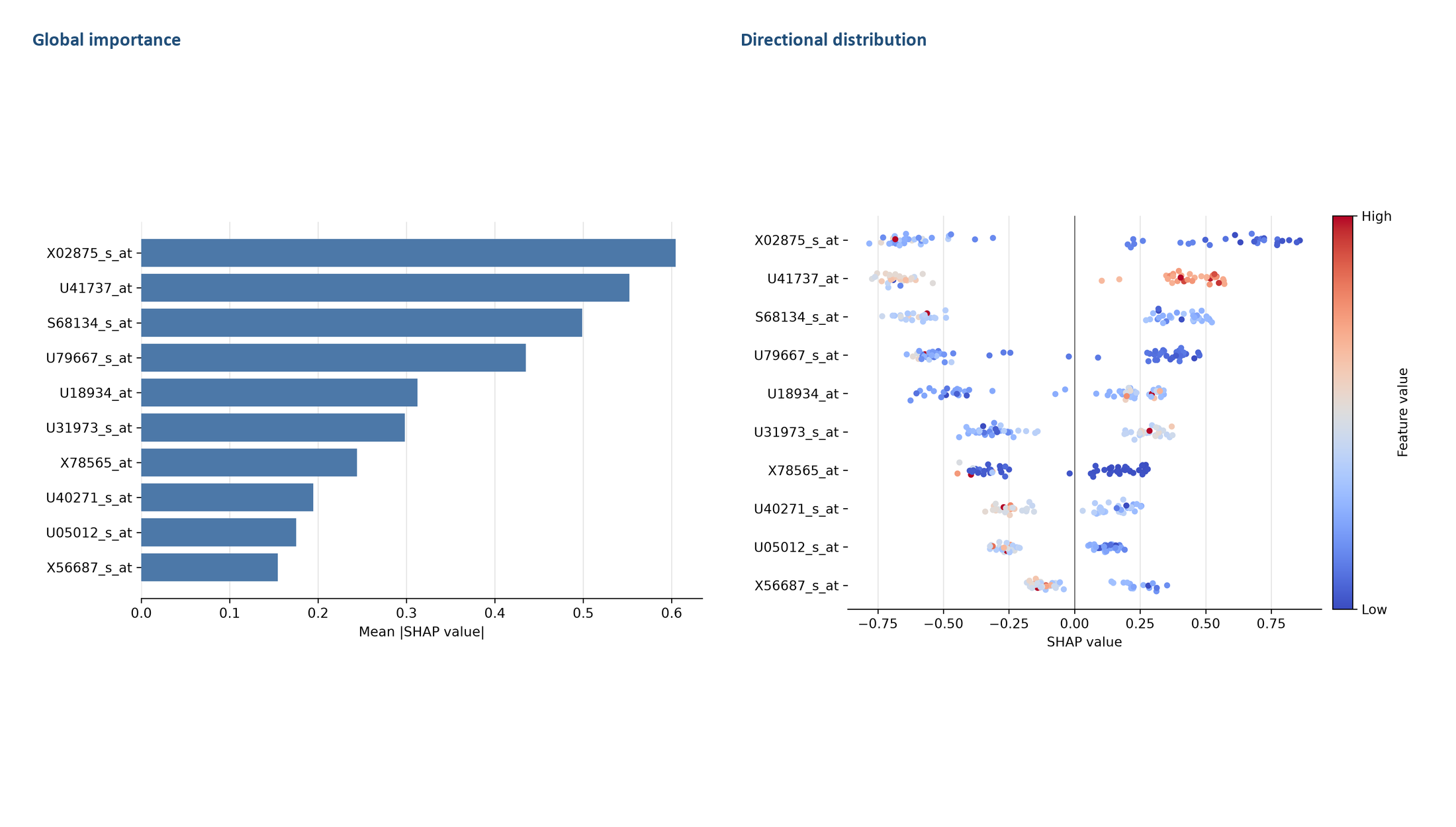

**Figure S2B.** SHAP summary for Binary BrainTumor. The left panel ranks features by mean absolute SHAP value. The right panel shows the distribution and direction of feature contributions across samples; red indicates higher feature values and blue indicates lower values. SHAP values indicate predictive attribution, not causality.

**Figure S2C**

Binary BreastCancer

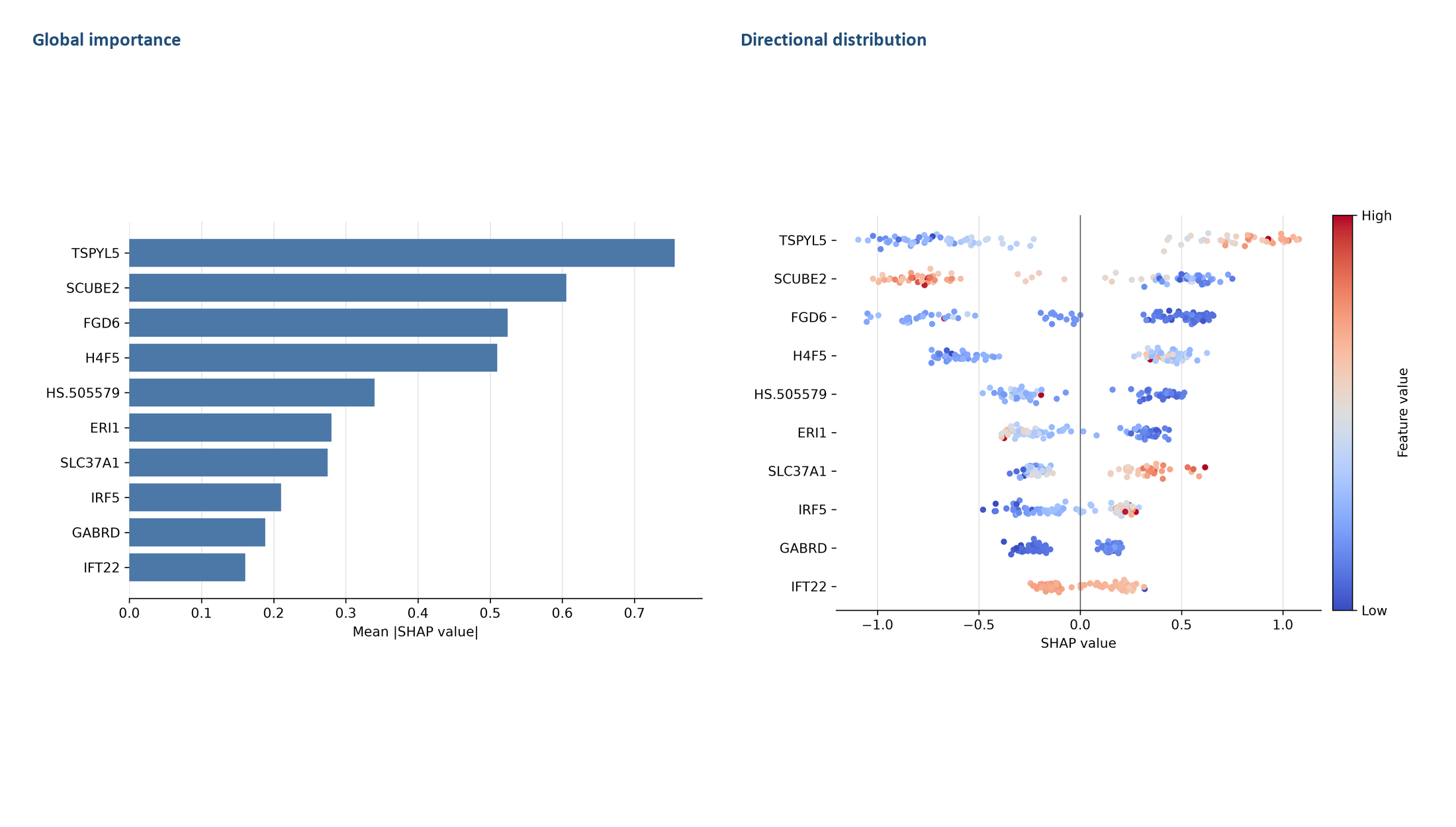

**Figure S2C.** SHAP summary for Binary BreastCancer. The left panel ranks features by mean absolute SHAP value. The right panel shows the distribution and direction of feature contributions across samples; red indicates higher feature values and blue indicates lower values. SHAP values indicate predictive attribution, not causality.

**Figure S2D**

Binary ColonTumor

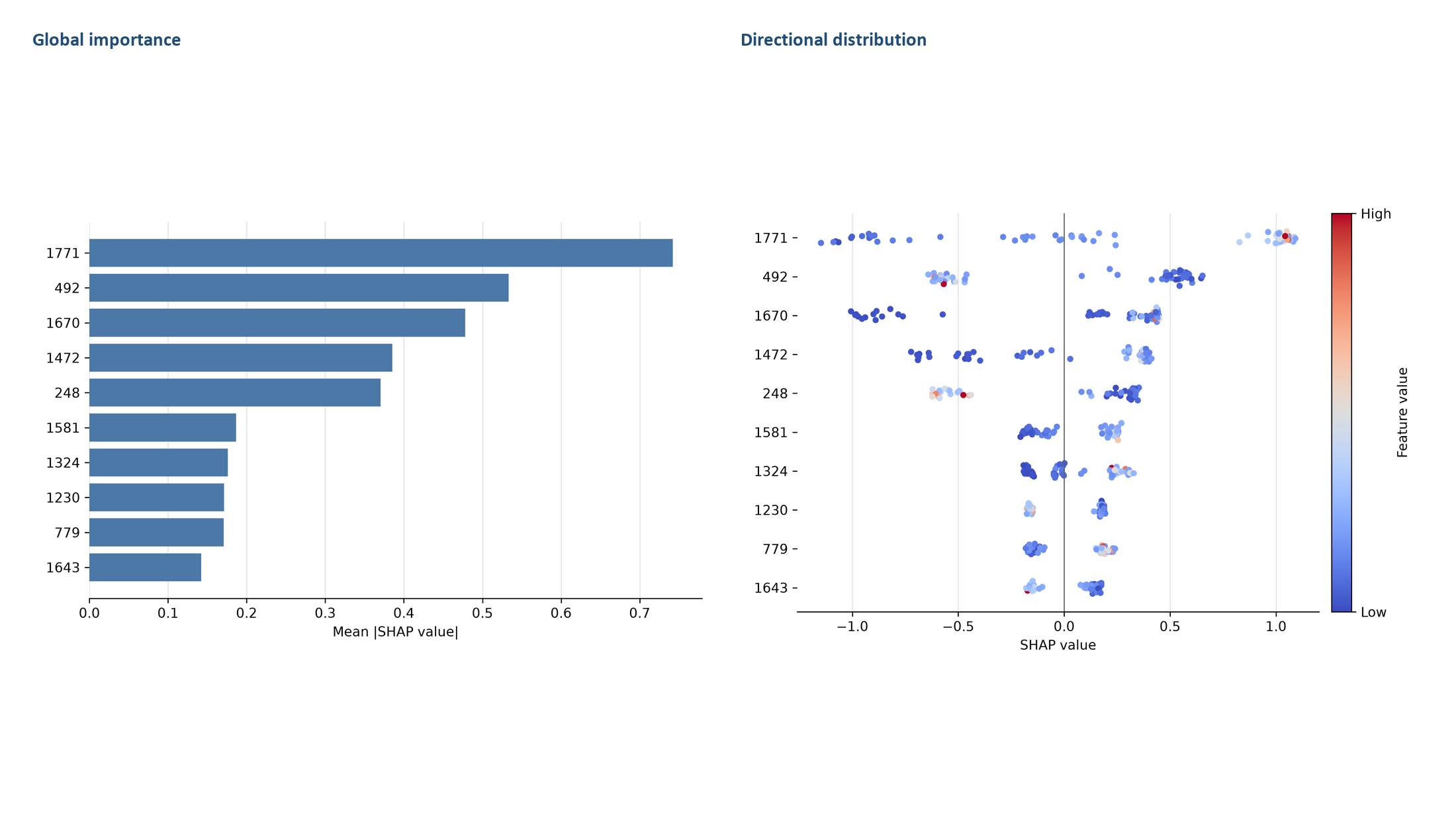

**Figure S2D.** SHAP summary for Binary ColonTumor. The left panel ranks features by mean absolute SHAP value. The right panel shows the distribution and direction of feature contributions across samples; red indicates higher feature values and blue indicates lower values. SHAP values indicate predictive attribution, not causality.

**Figure S2E**

Binary Gastric

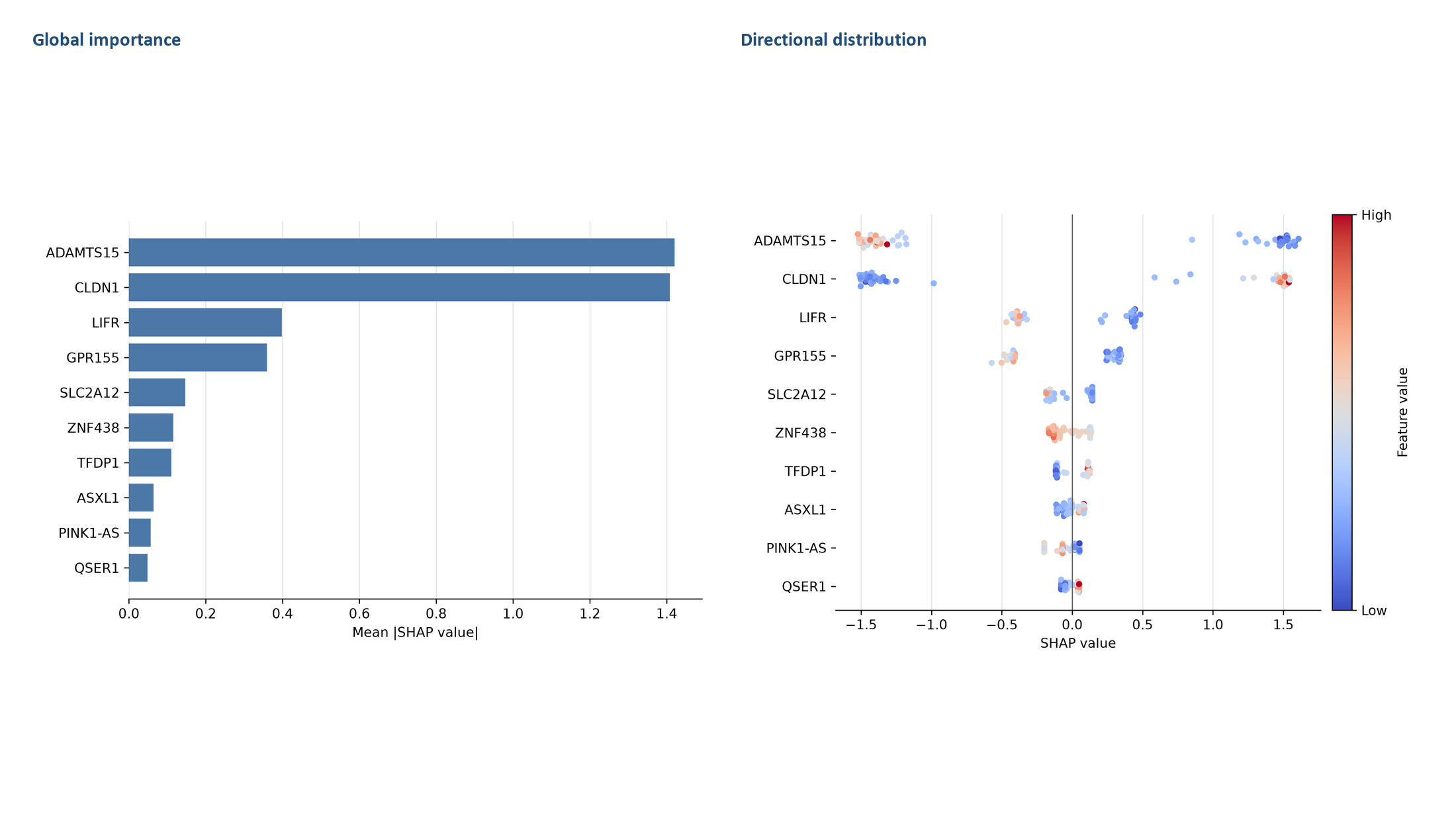

**Figure S2E.** SHAP summary for Binary Gastric. The left panel ranks features by mean absolute SHAP value. The right panel shows the distribution and direction of feature contributions across samples; red indicates higher feature values and blue indicates lower values. SHAP values indicate predictive attribution, not causality.

**Figure S2F**

Binary Leukemia

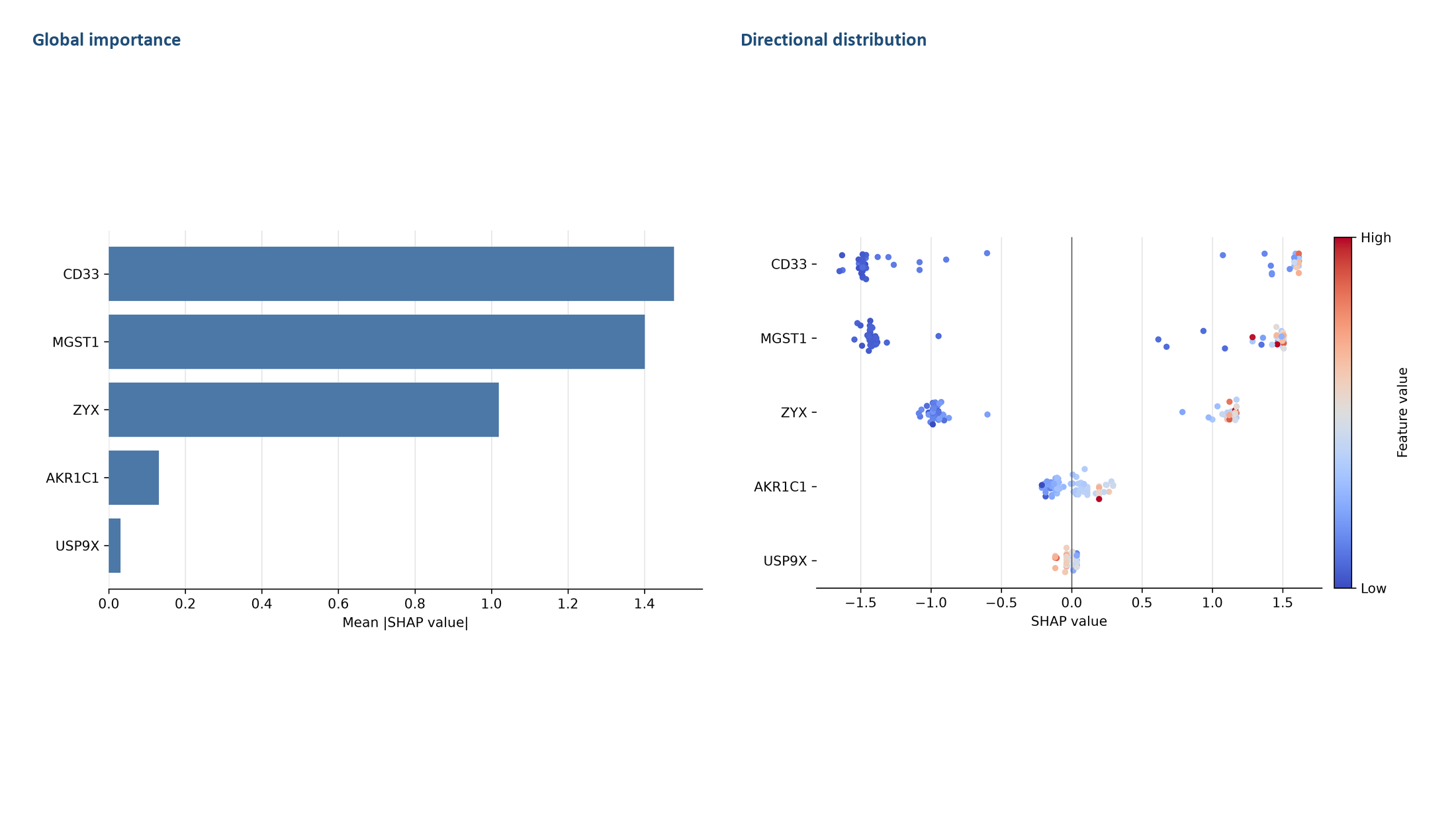

**Figure S2F.** SHAP summary for Binary Leukemia. The left panel ranks features by mean absolute SHAP value. The right panel shows the distribution and direction of feature contributions across samples; red indicates higher feature values and blue indicates lower values. SHAP values indicate predictive attribution, not causality.

**Figure S2G**

Binary Lung

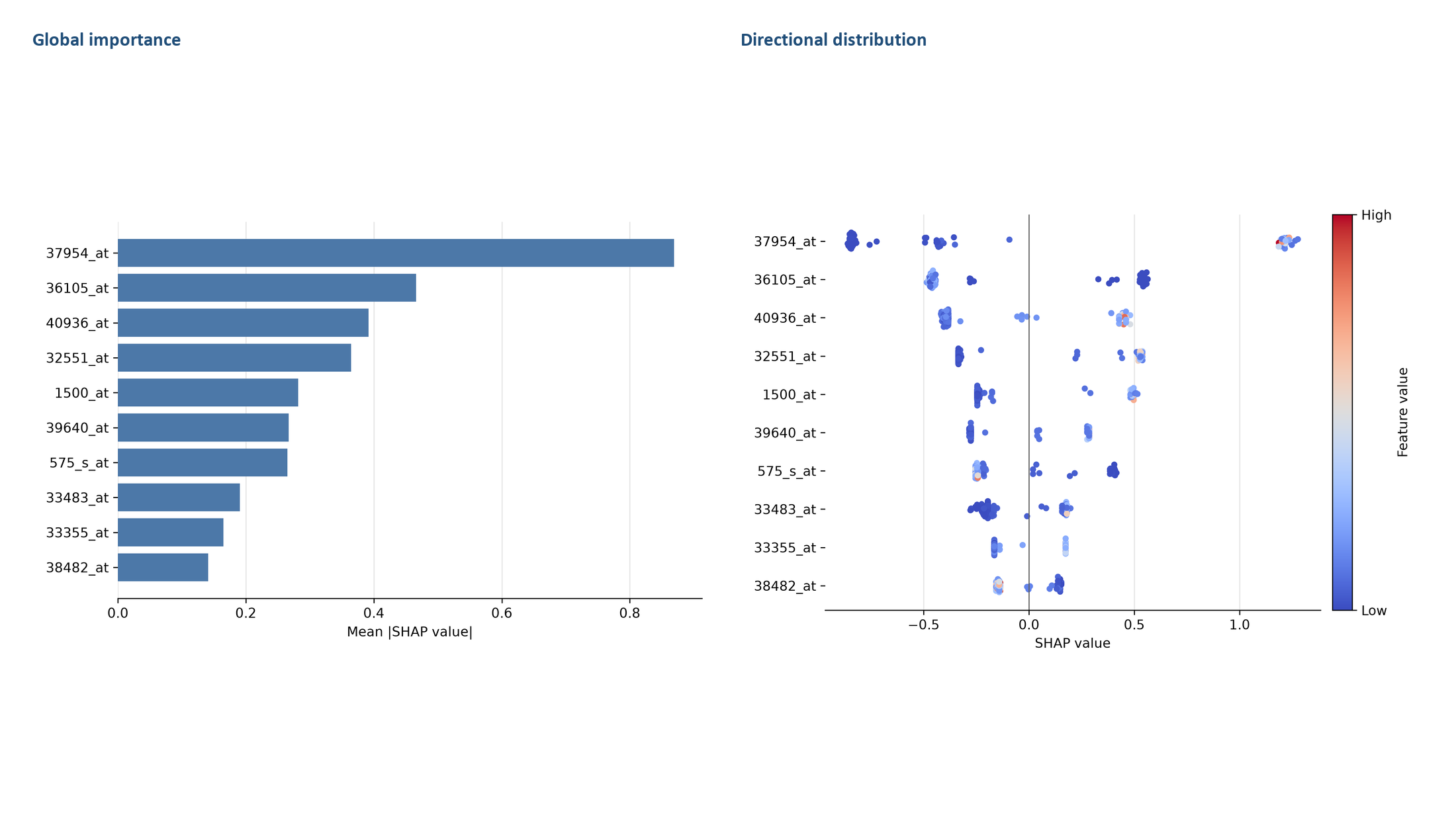

**Figure S2G.** SHAP summary for Binary Lung. The left panel ranks features by mean absolute SHAP value. The right panel shows the distribution and direction of feature contributions across samples; red indicates higher feature values and blue indicates lower values. SHAP values indicate predictive attribution, not causality.

**Figure S2H**

Binary Lymphoma

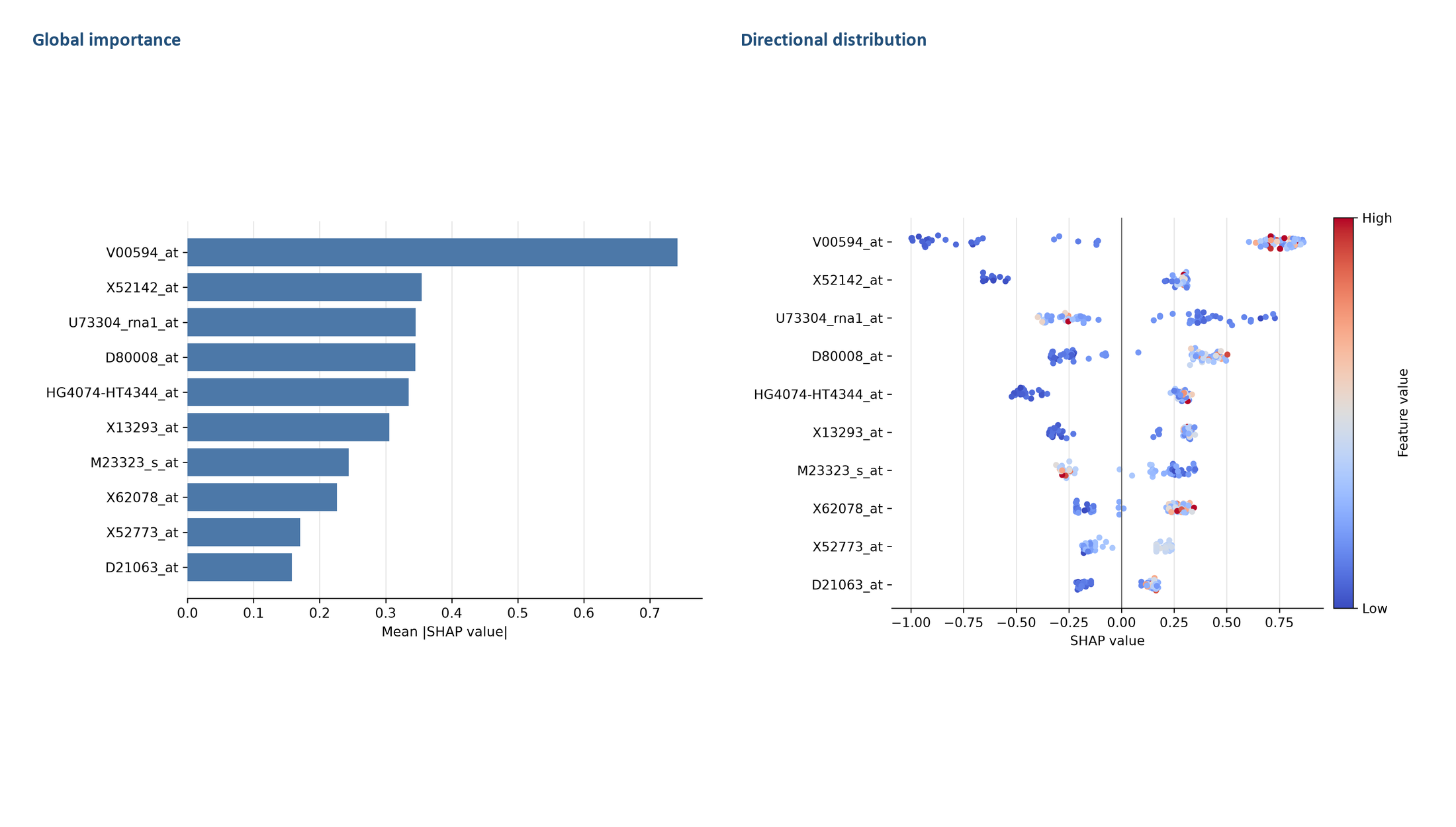

**Figure S2H.** SHAP summary for Binary Lymphoma. The left panel ranks features by mean absolute SHAP value. The right panel shows the distribution and direction of feature contributions across samples; red indicates higher feature values and blue indicates lower values. SHAP values indicate predictive attribution, not causality.

**Figure S2I**

Binary Myeloma

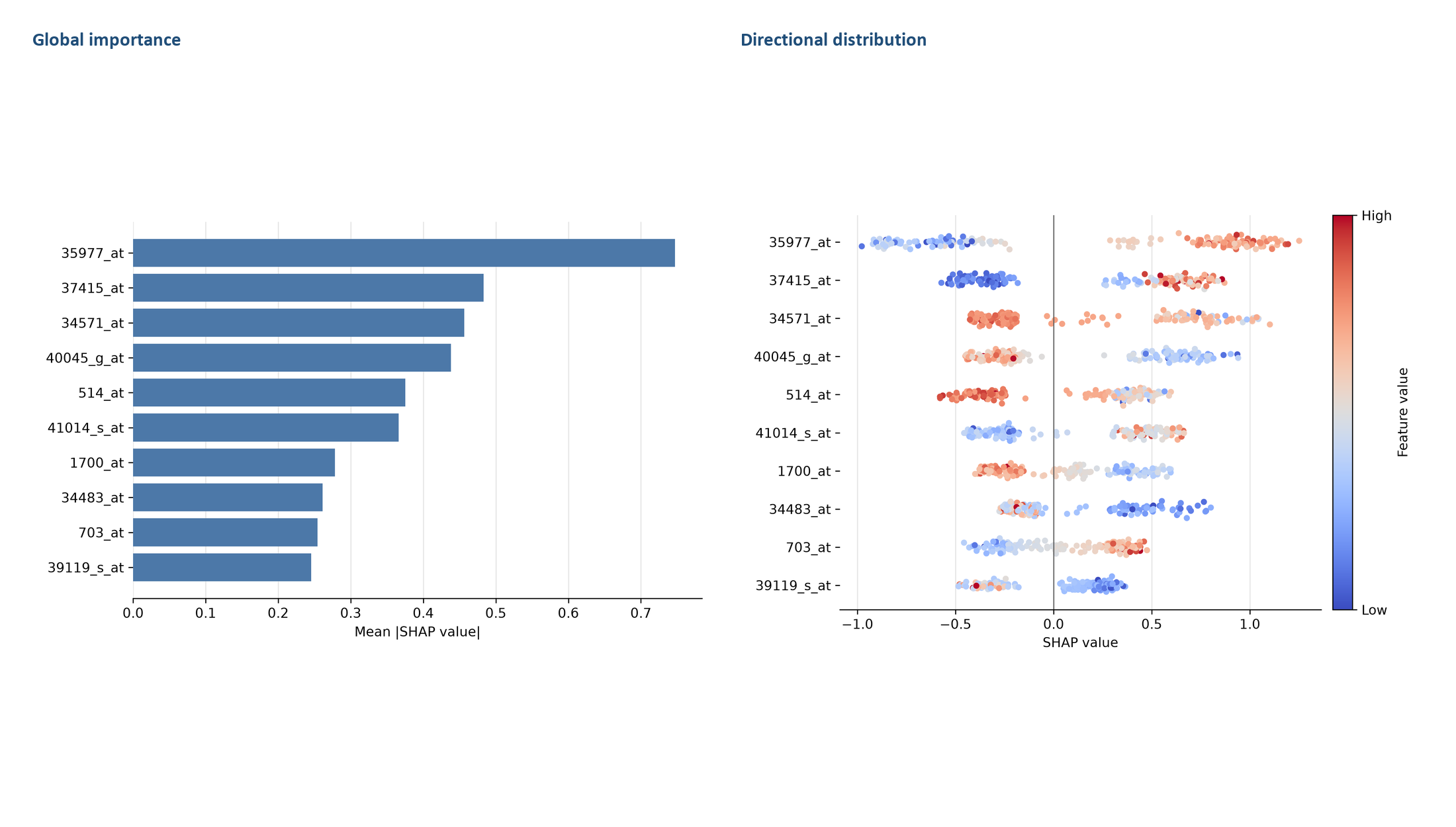

**Figure S2I.** SHAP summary for Binary Myeloma. The left panel ranks features by mean absolute SHAP value. The right panel shows the distribution and direction of feature contributions across samples; red indicates higher feature values and blue indicates lower values. SHAP values indicate predictive attribution, not causality.

**Figure S2J**

Binary OvarianCancer

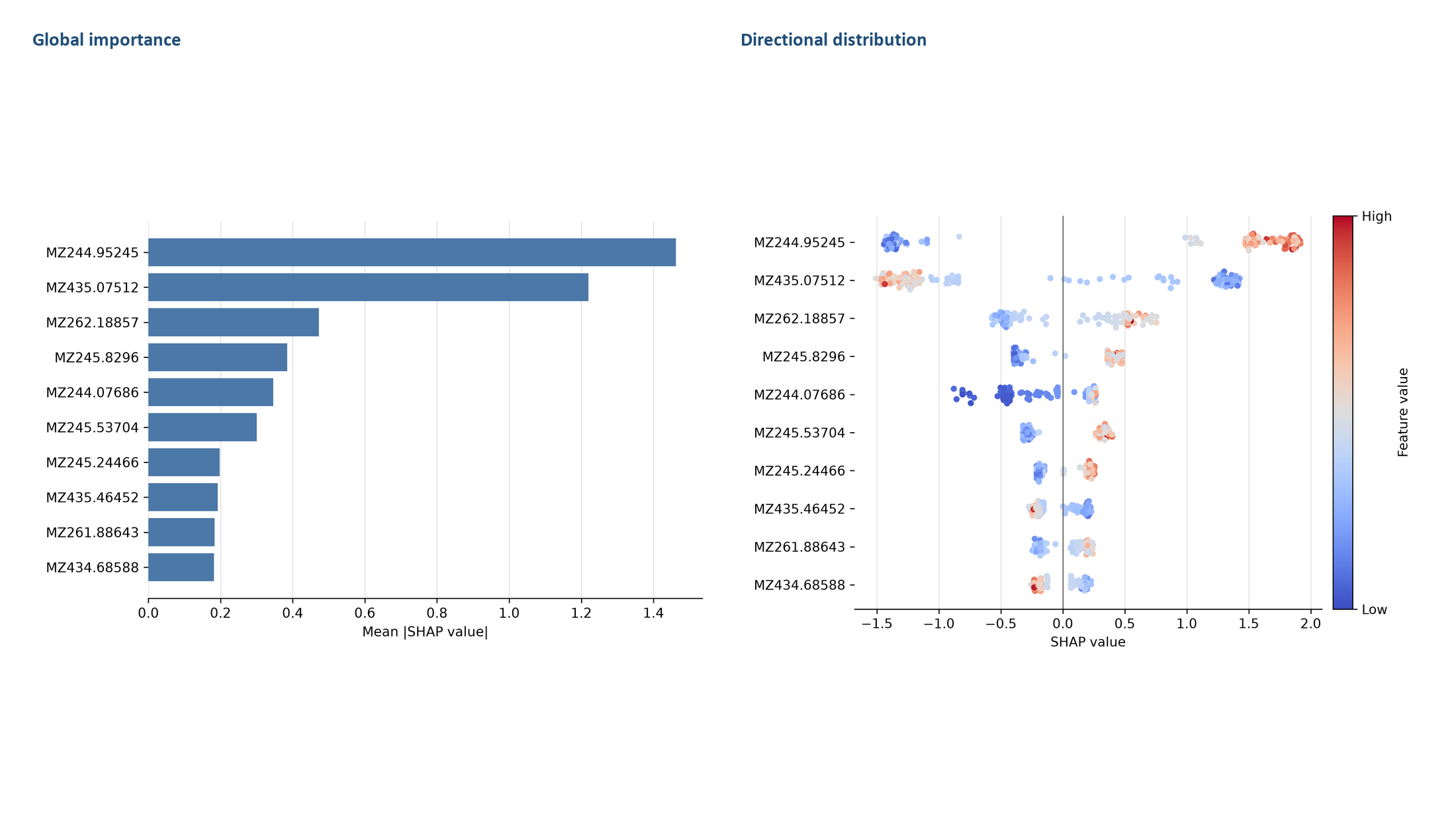

**Figure S2J.** SHAP summary for Binary OvarianCancer. The left panel ranks features by mean absolute SHAP value. The right panel shows the distribution and direction of feature contributions across samples; red indicates higher feature values and blue indicates lower values. SHAP values indicate predictive attribution, not causality.

**Figure S2K**

Binary Prostate

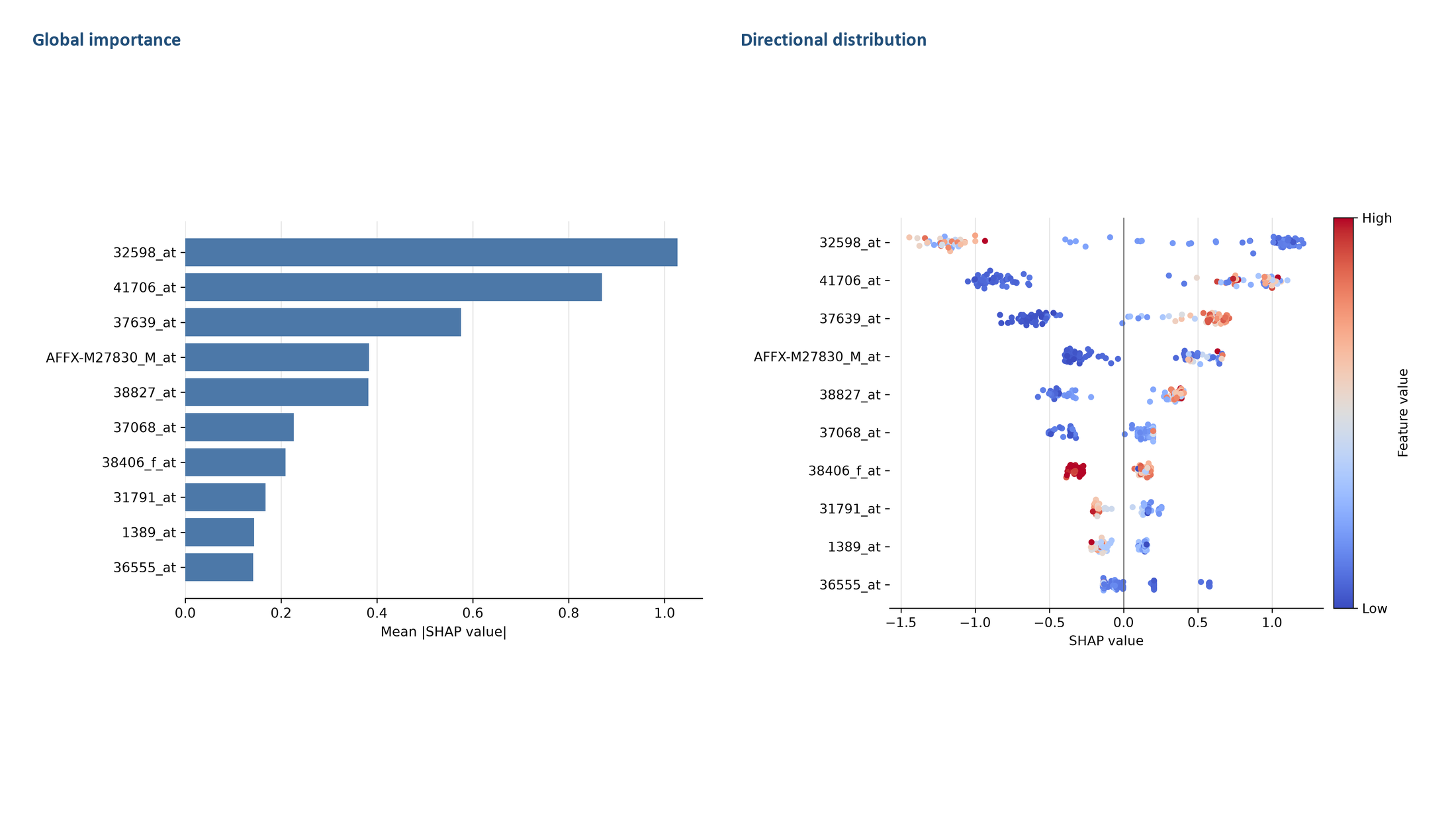

**Figure S2K.** SHAP summary for Binary Prostate. The left panel ranks features by mean absolute SHAP value. The right panel shows the distribution and direction of feature contributions across samples; red indicates higher feature values and blue indicates lower values. SHAP values indicate predictive attribution, not causality.

**Figure S3A**

Multiclass BrainCancer — overview

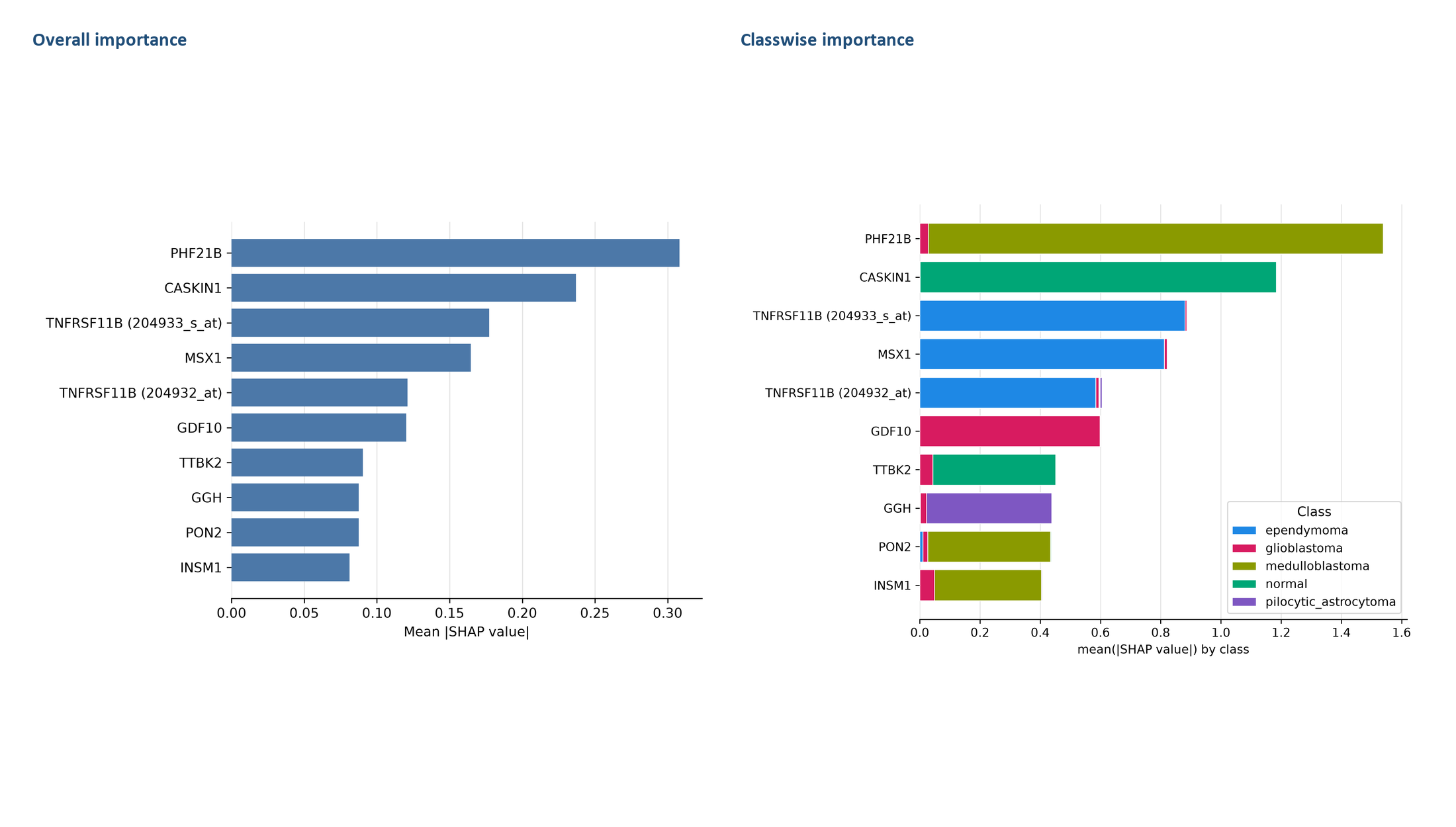

**Figure S3A.** Multiclass SHAP summary for Multiclass BrainCancer. Overall and, where retained, classwise mean absolute SHAP summaries are shown. Attribution values quantify model contribution and do not establish biological causality.

**Figure S3A**

Multiclass BrainCancer — class-specific panels 1/2

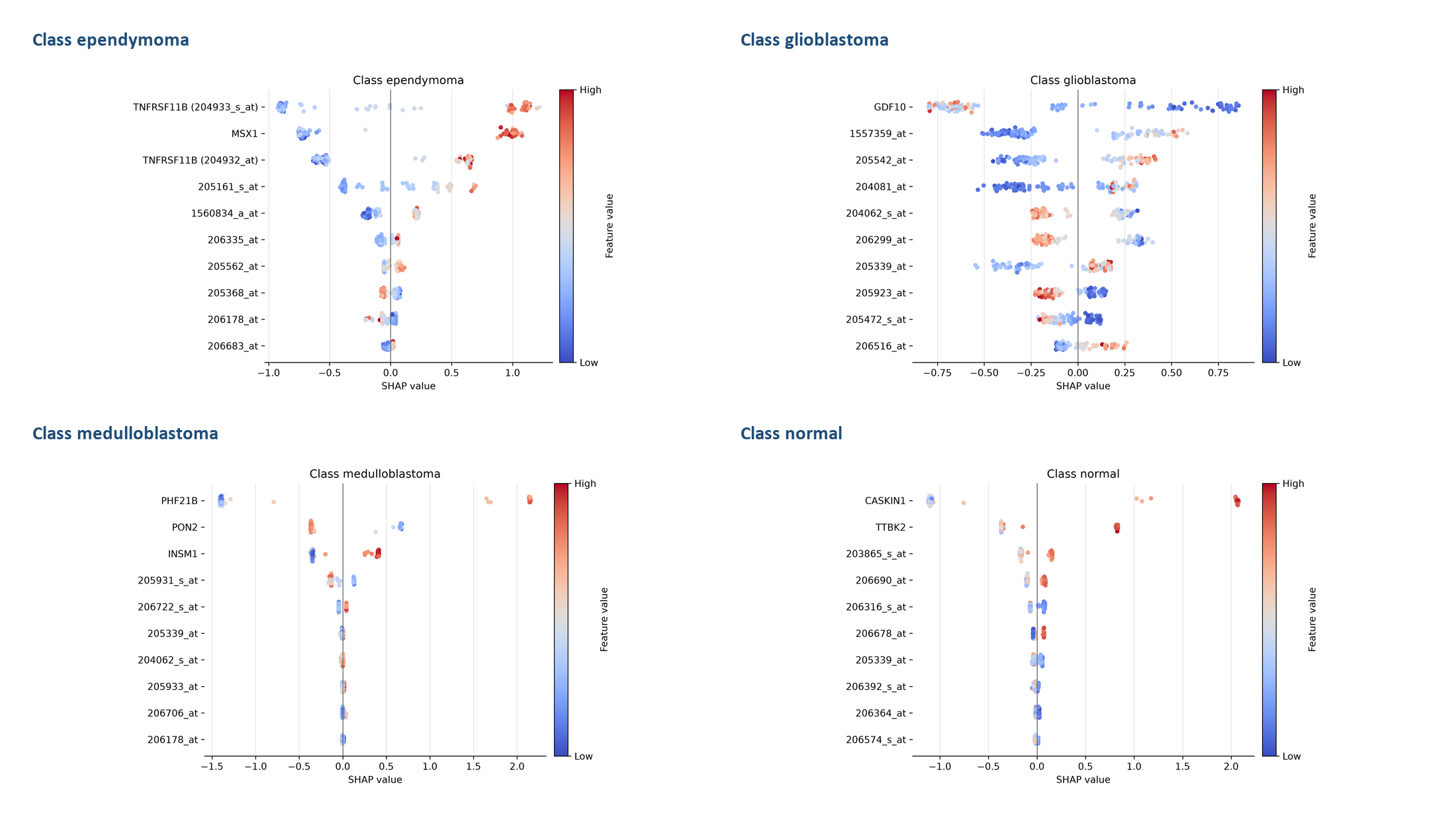

**Figure S3A (continued).** Class-specific SHAP distributions for Multiclass BrainCancer. Each panel shows the magnitude and direction of feature contributions for the indicated class; red denotes higher feature values and blue denotes lower values.

**Figure S3A**

Multiclass BrainCancer — class-specific panels 2/2

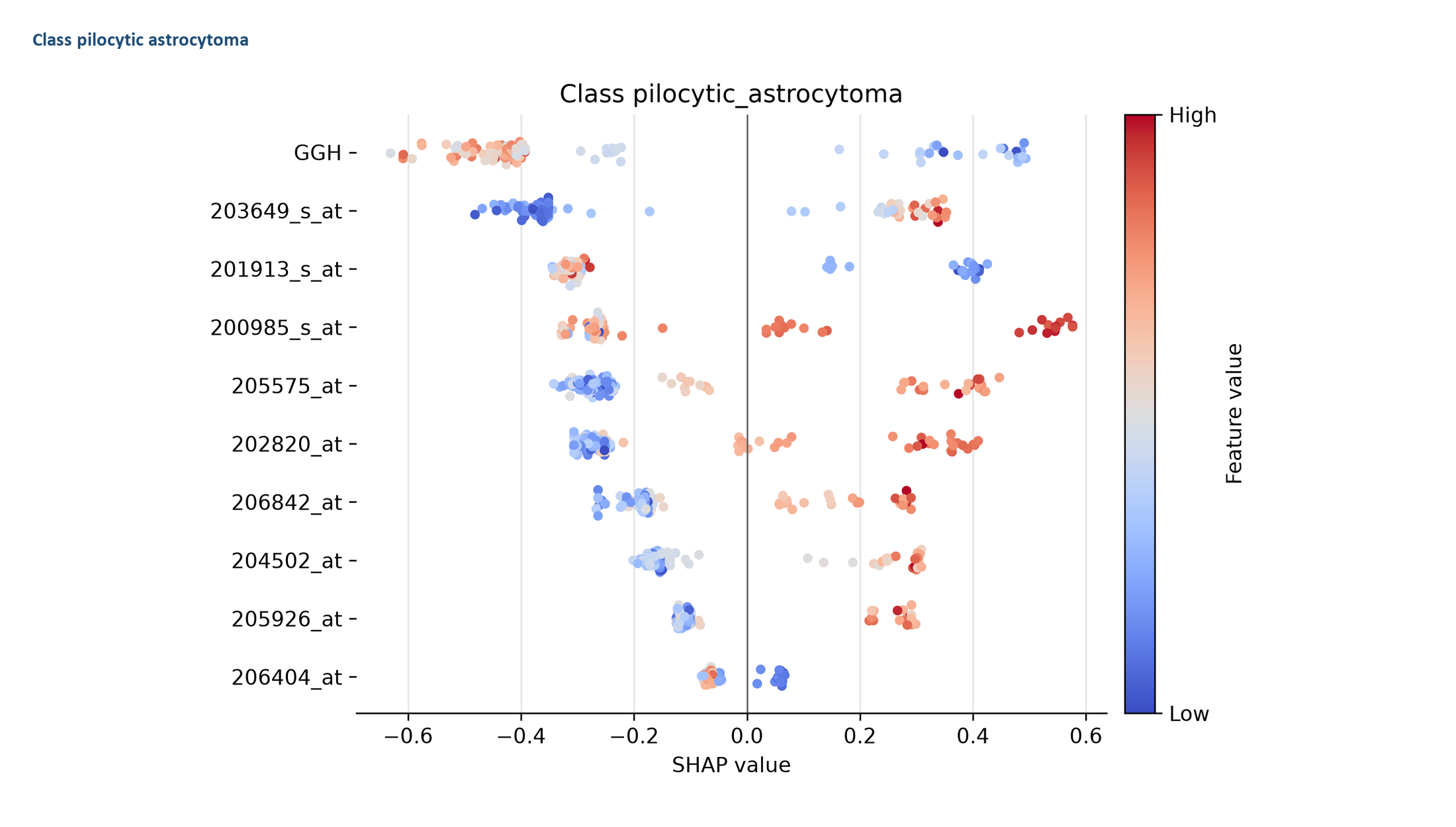

**Figure S3A (continued).** Class-specific SHAP distributions for Multiclass BrainCancer. Each panel shows the magnitude and direction of feature contributions for the indicated class; red denotes higher feature values and blue denotes lower values.

**Figure S3B**

Multiclass Crohns — overview

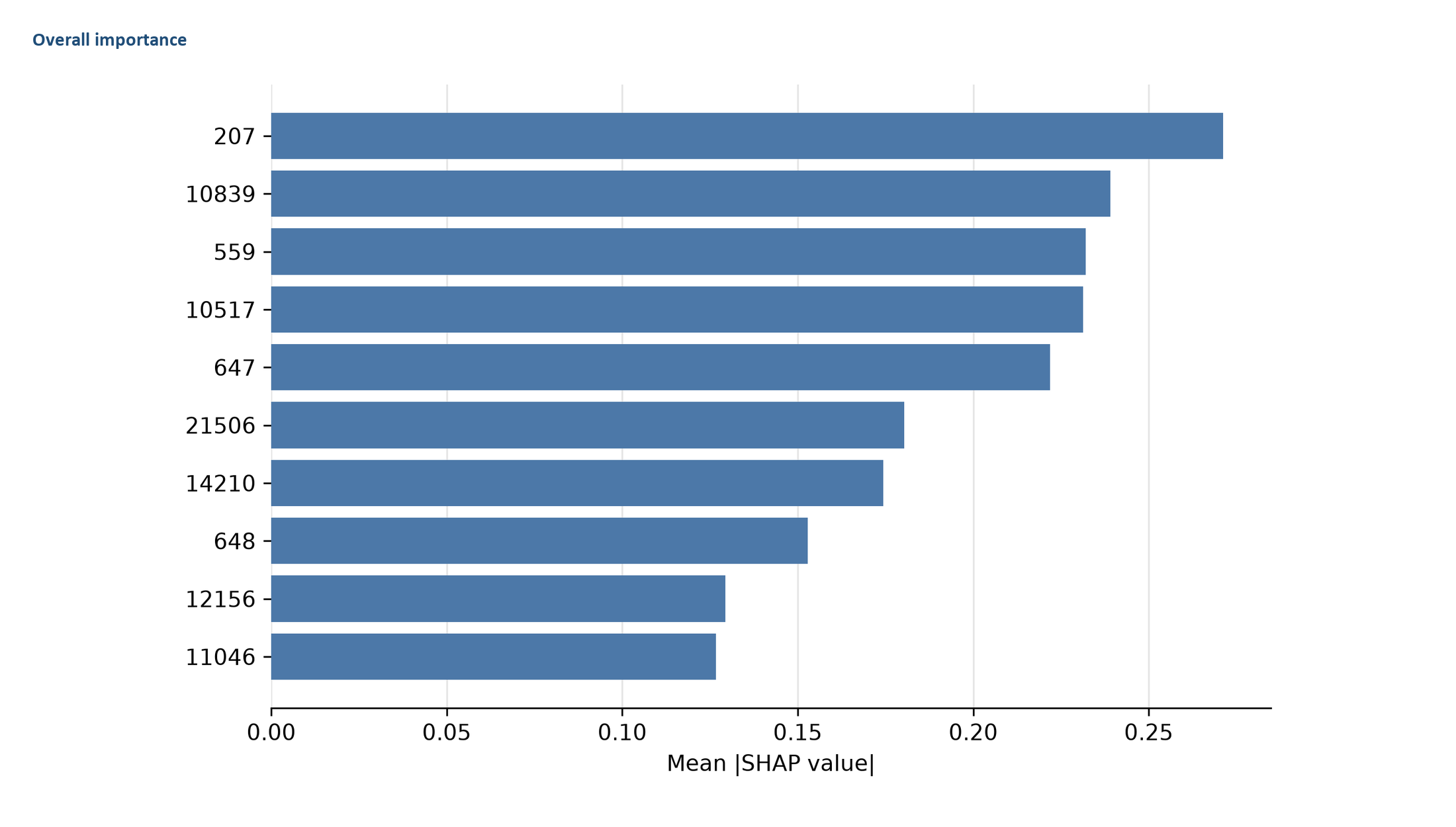

**Figure S3B.** Multiclass SHAP summary for Multiclass Crohns. Overall and, where retained, classwise mean absolute SHAP summaries are shown. Attribution values quantify model contribution and do not establish biological causality.

**Figure S3B**

Multiclass Crohns — class-specific panels 1/1

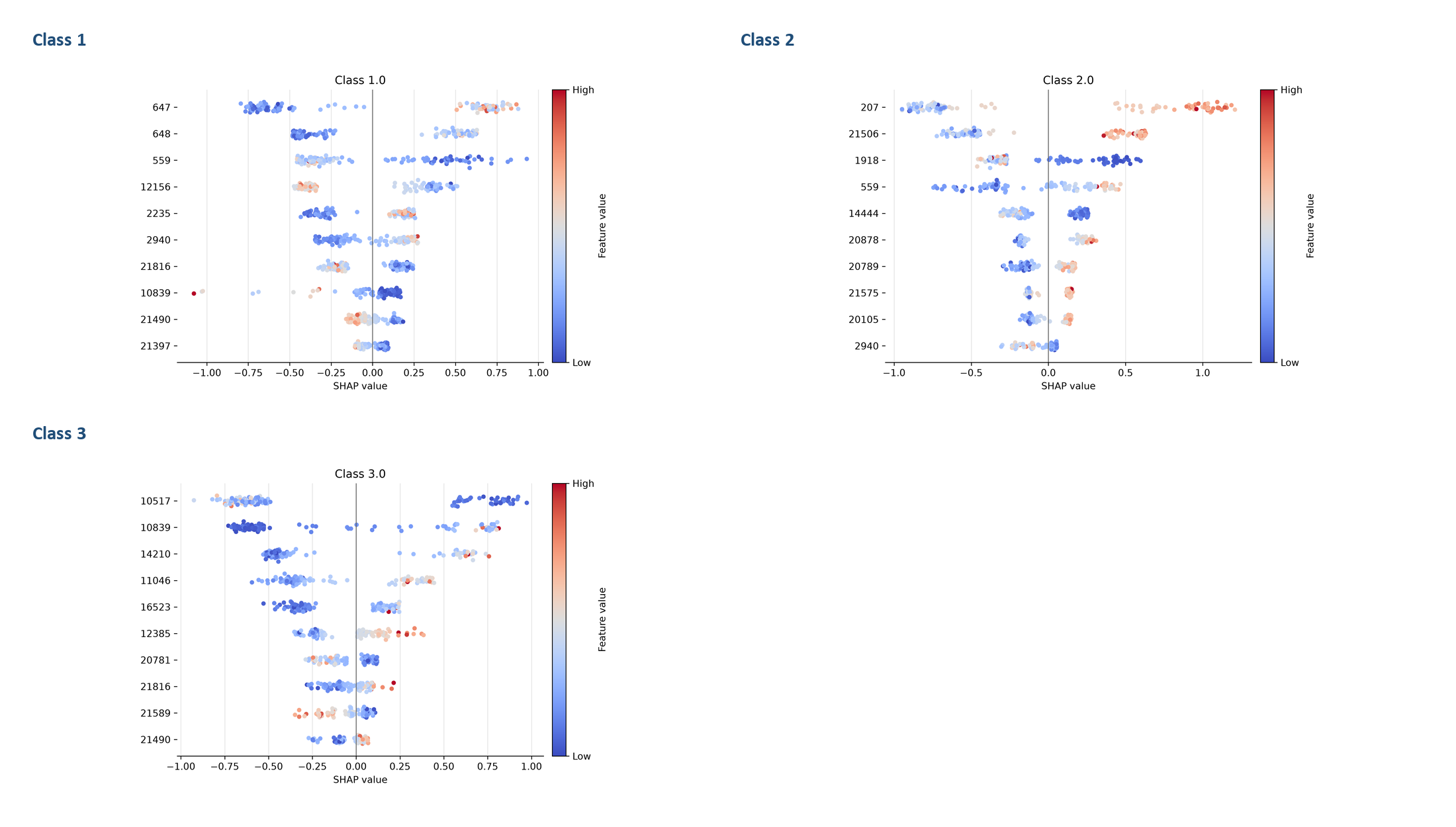

**Figure S3B (continued).** Class-specific SHAP distributions for Multiclass Crohns. Each panel shows the magnitude and direction of feature contributions for the indicated class; red denotes higher feature values and blue denotes lower values.

**Figure S3C**

Multiclass EndometrialCancer — overview

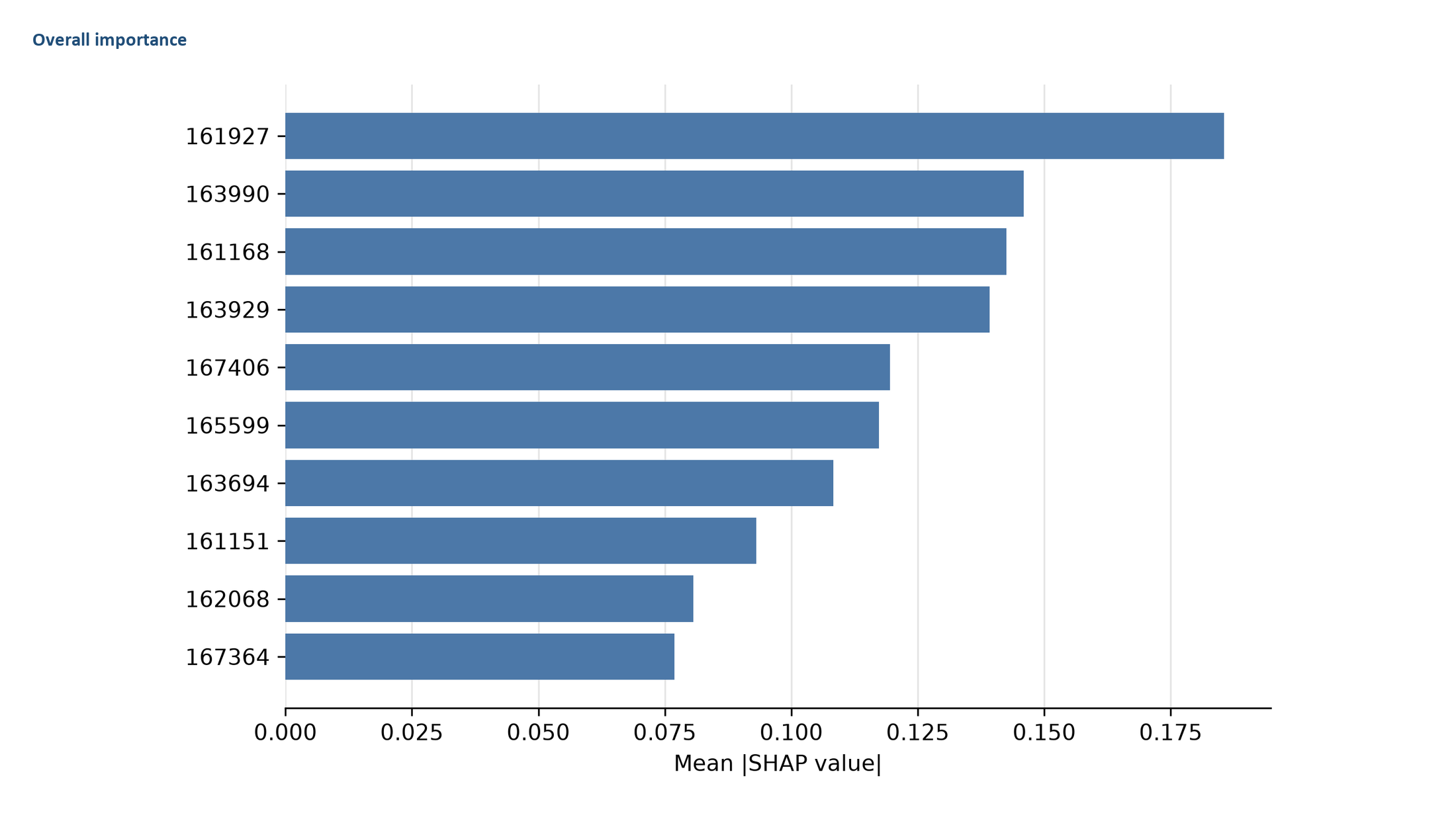

**Figure S3C.** Multiclass SHAP summary for Multiclass EndometrialCancer. Overall and, where retained, classwise mean absolute SHAP summaries are shown. Attribution values quantify model contribution and do not establish biological causality.

**Figure S3C**

Multiclass EndometrialCancer — class-specific panels 1/1

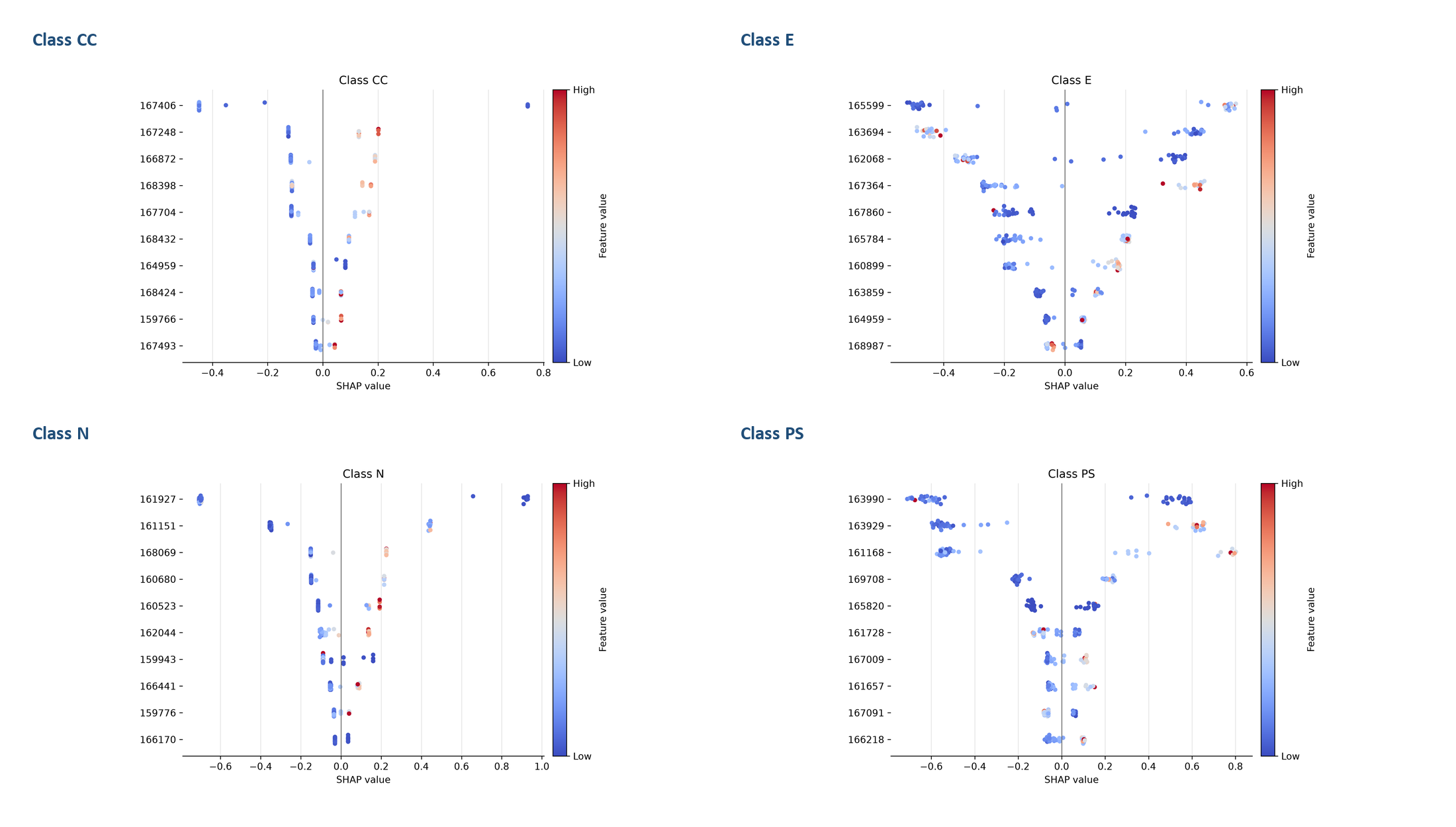

**Figure S3C (continued).** Class-specific SHAP distributions for Multiclass EndometrialCancer. Each panel shows the magnitude and direction of feature contributions for the indicated class; red denotes higher feature values and blue denotes lower values.

**Figure S3D**

Multiclass Glioma — overview

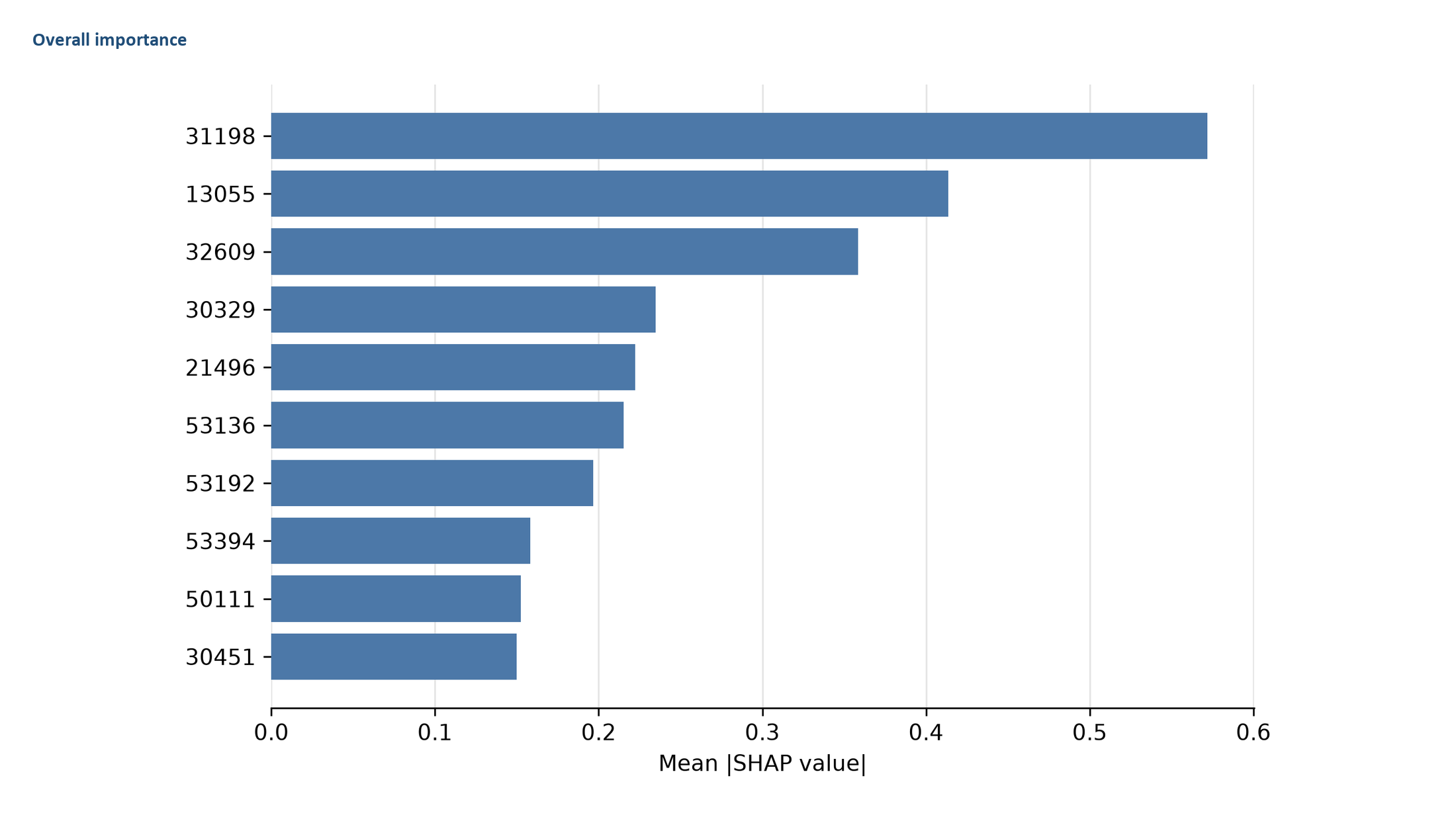

**Figure S3D.** Multiclass SHAP summary for Multiclass Glioma. Overall and, where retained, classwise mean absolute SHAP summaries are shown. Attribution values quantify model contribution and do not establish biological causality.

**Figure S3D**

Multiclass Glioma — class-specific panels 1/1

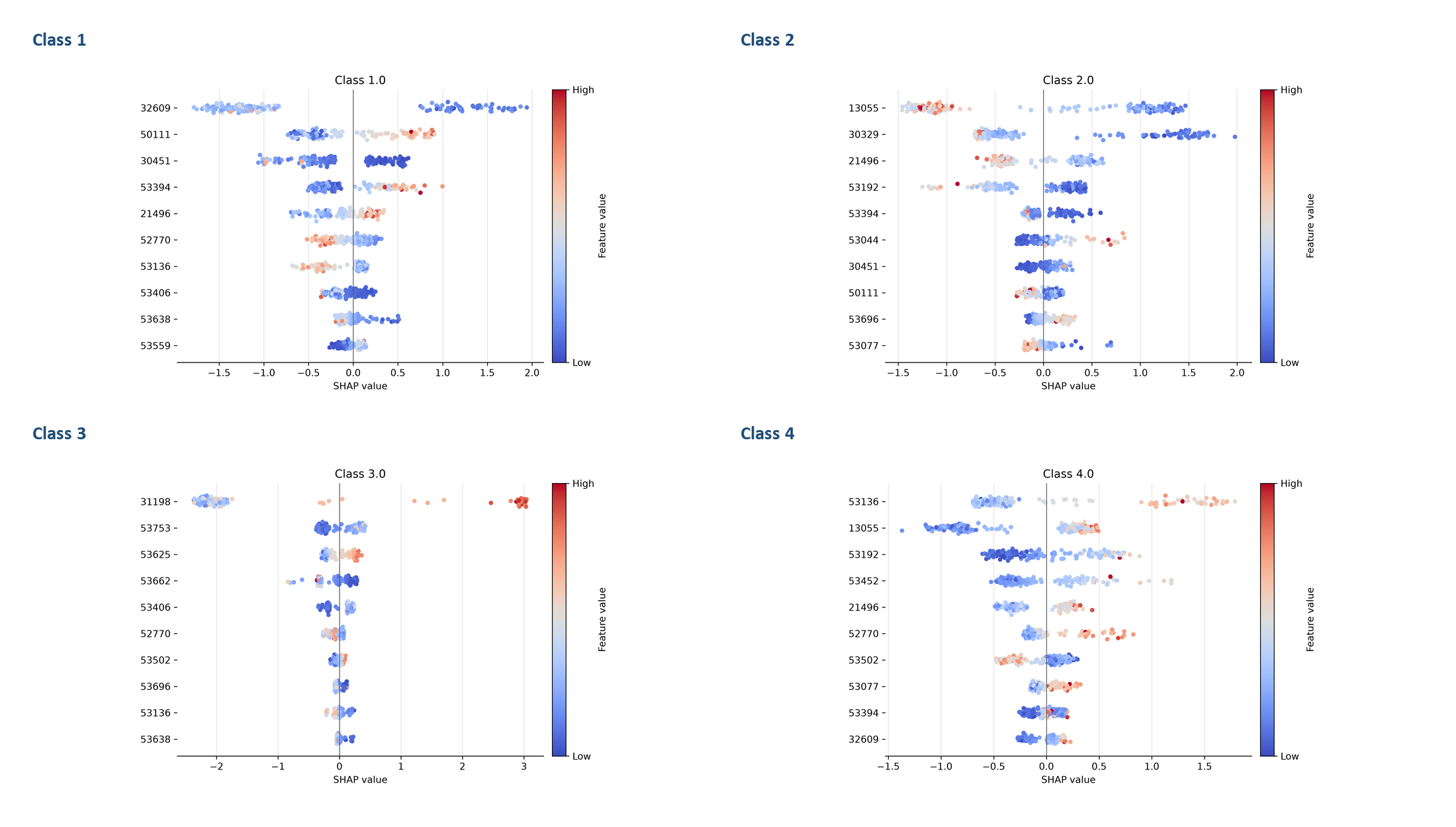

**Figure S3D (continued).** Class-specific SHAP distributions for Multiclass Glioma. Each panel shows the magnitude and direction of feature contributions for the indicated class; red denotes higher feature values and blue denotes lower values.

**Figure S3E**

Multiclass Leukemia 3-class — overview

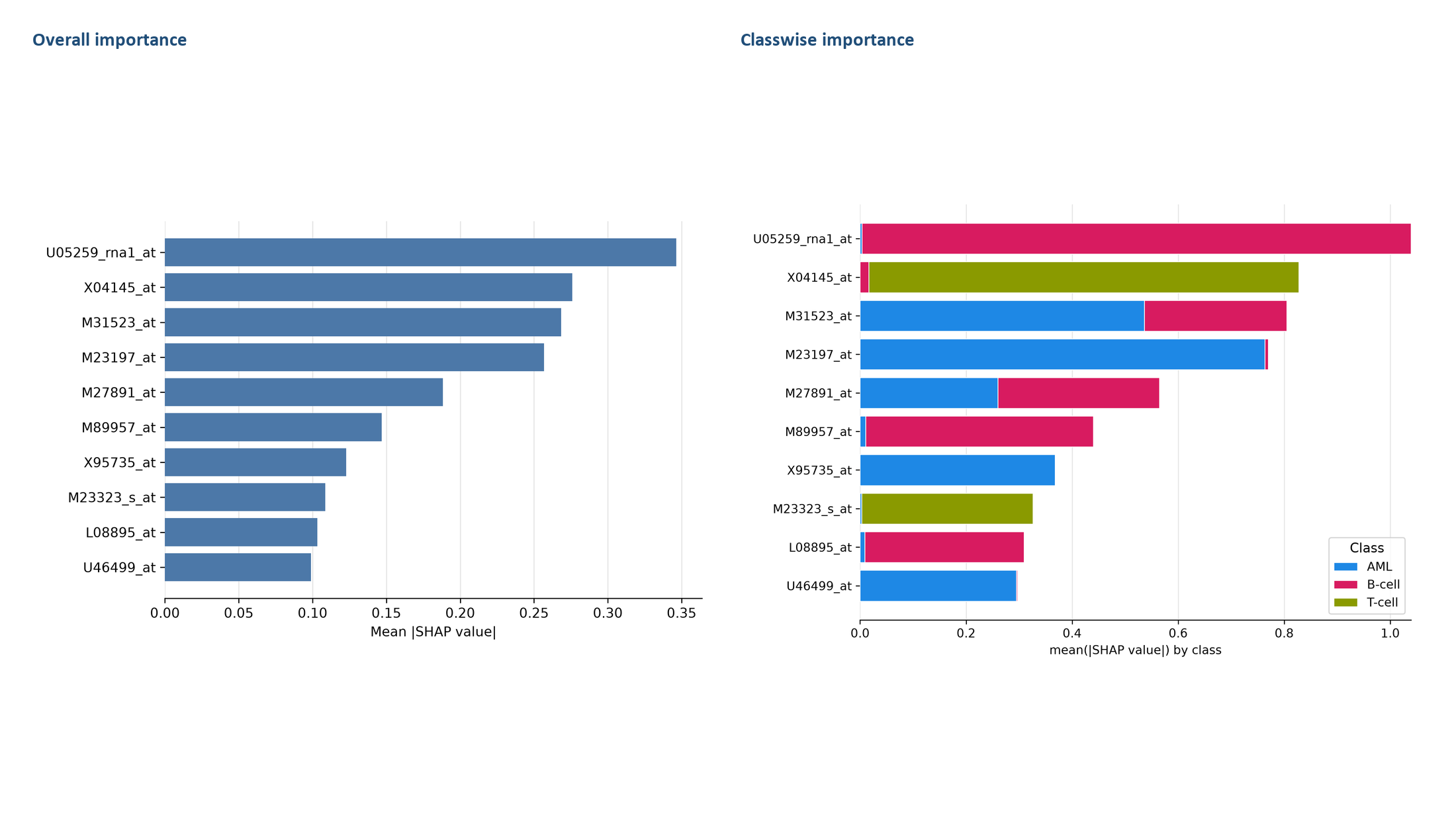

**Figure S3E.** Multiclass SHAP summary for Multiclass Leukemia 3. Overall and, where retained, classwise mean absolute SHAP summaries are shown. Attribution values quantify model contribution and do not establish biological causality.

**Figure S3E**

Multiclass Leukemia 3-class — class-specific panels 1/1

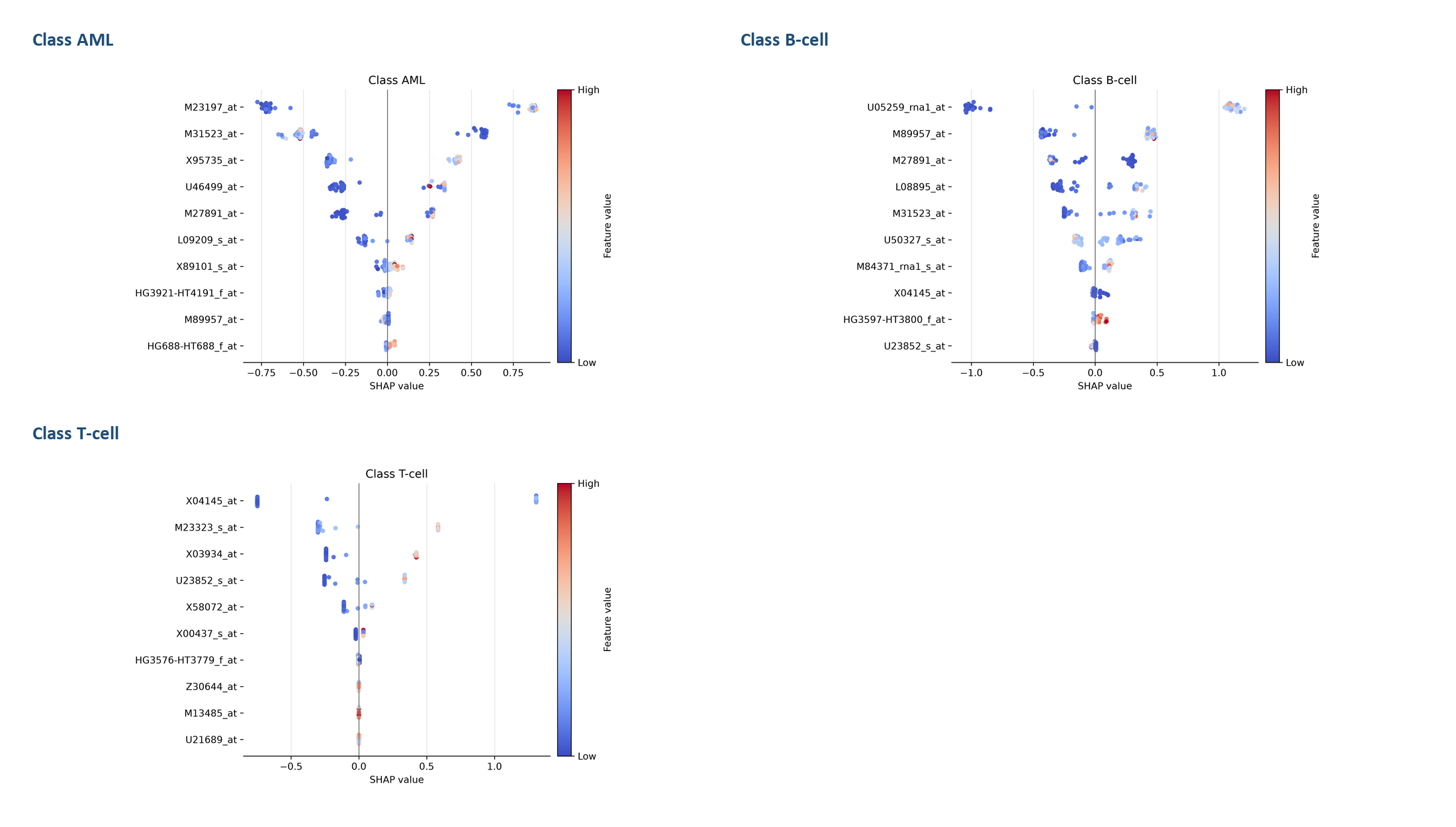

**Figure S3E (continued).** Class-specific SHAP distributions for Multiclass Leukemia 3. Each panel shows the magnitude and direction of feature contributions for the indicated class; red denotes higher feature values and blue denotes lower values.

**Figure S3F**

Multiclass Leukemia 4-class — overview

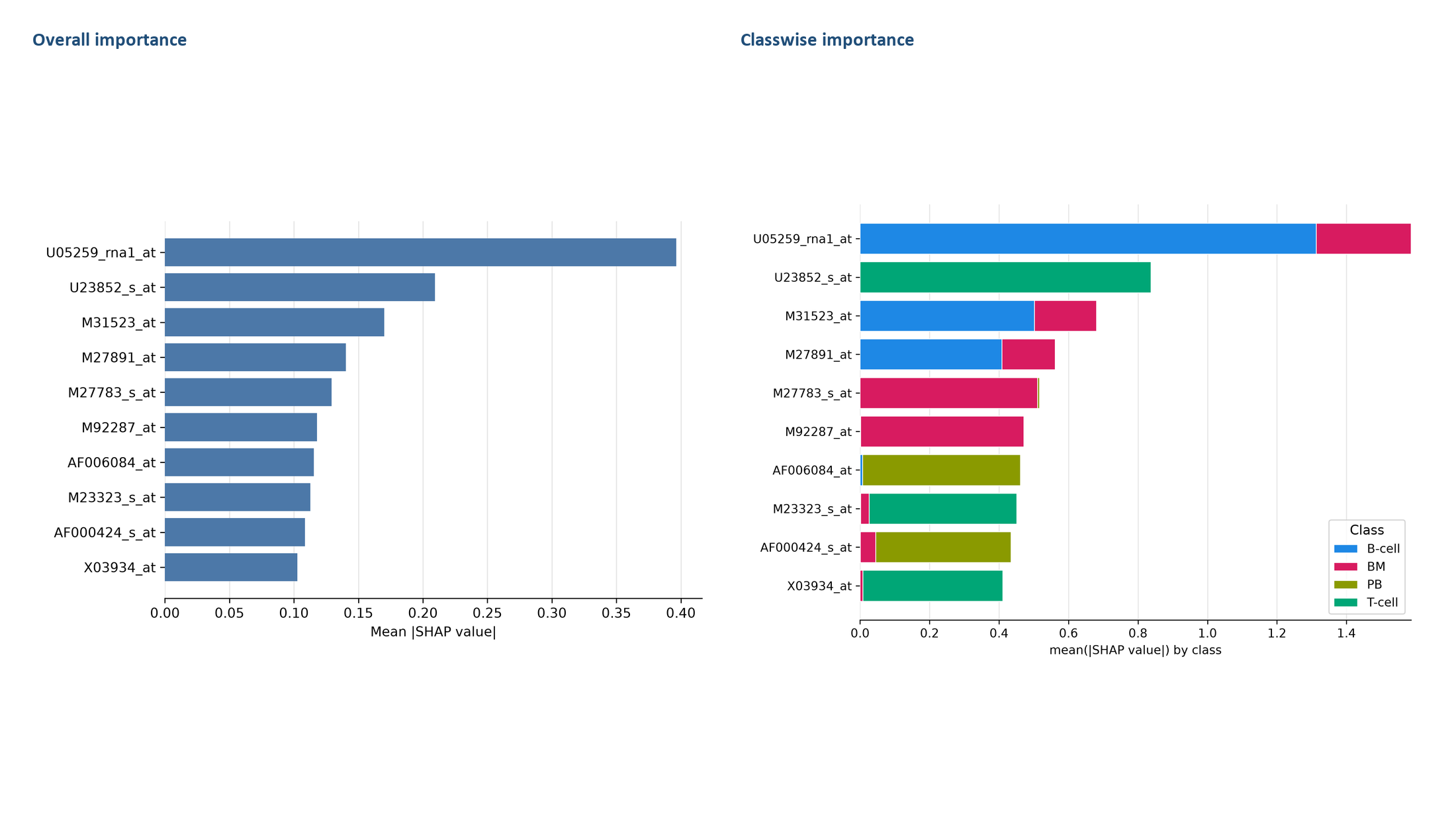

**Figure S3F.** Multiclass SHAP summary for Multiclass Leukemia 4. Overall and, where retained, classwise mean absolute SHAP summaries are shown. Attribution values quantify model contribution and do not establish biological causality.

**Figure S3F**

Multiclass Leukemia 4-class — class-specific panels 1/1

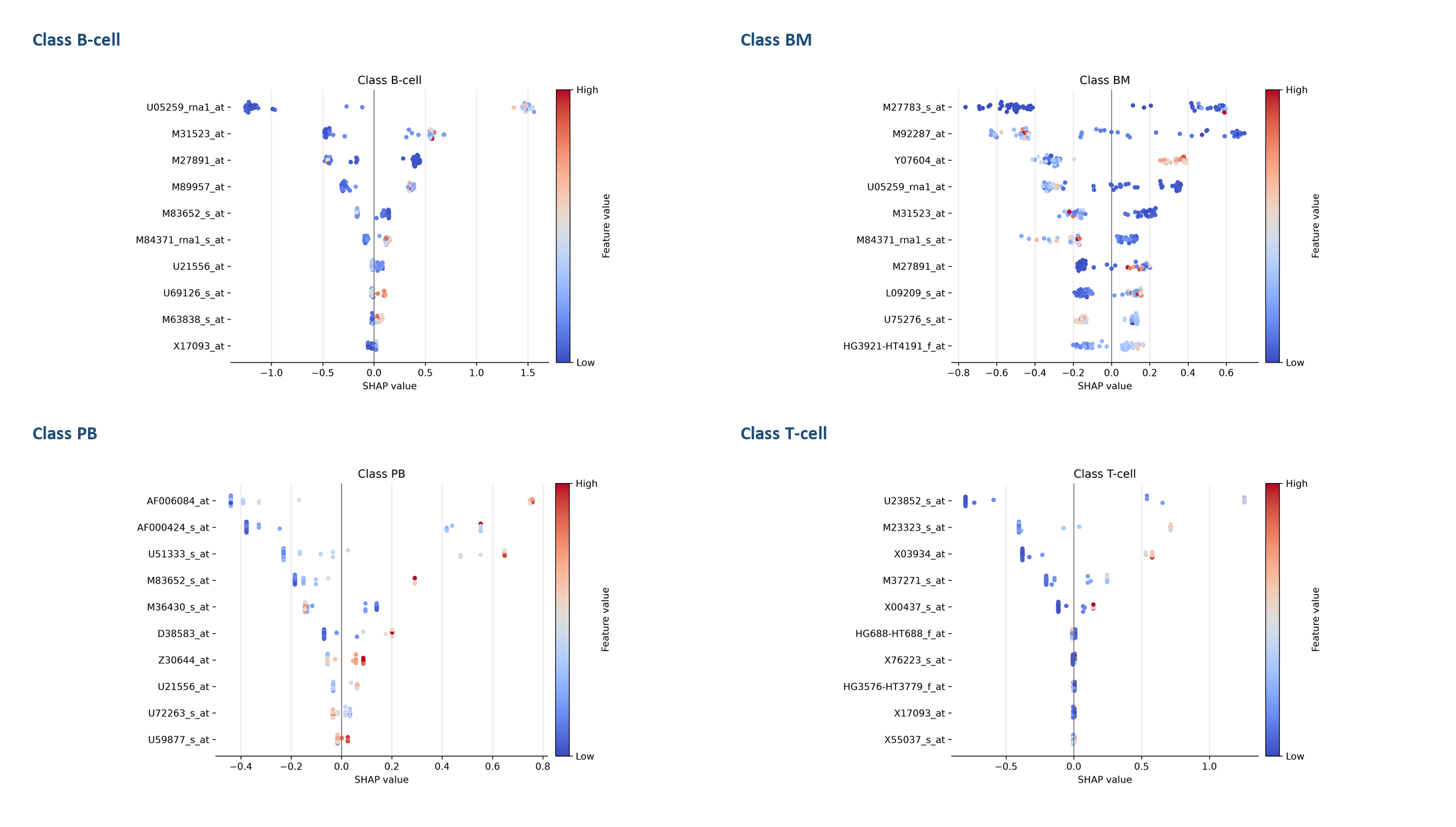

**Figure S3F (continued).** Class-specific SHAP distributions for Multiclass Leukemia 4. Each panel shows the magnitude and direction of feature contributions for the indicated class; red denotes higher feature values and blue denotes lower values.

**Figure S3G**

Multiclass LungCancer — overview

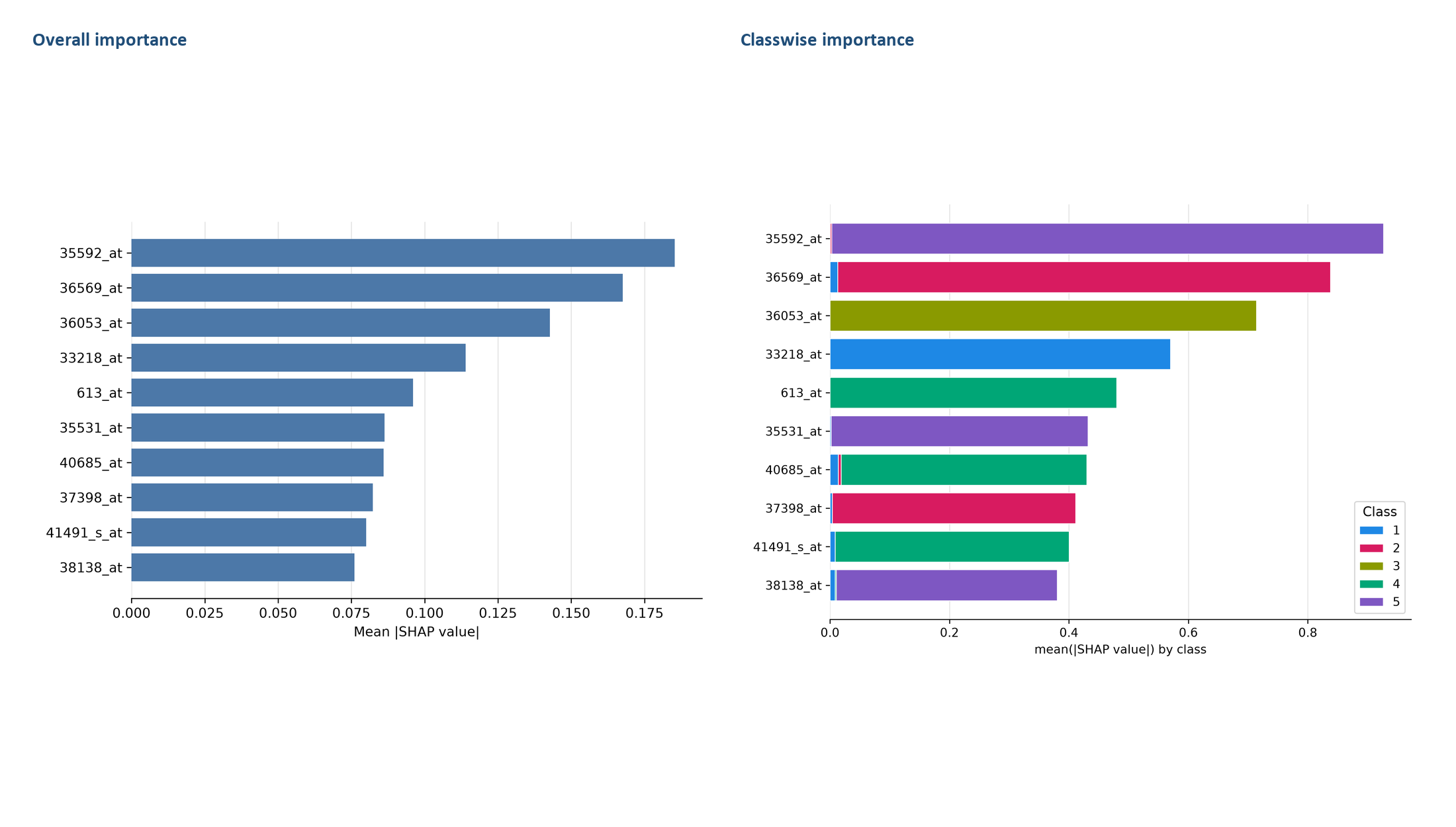

**Figure S3G.** Multiclass SHAP summary for Multiclass LungCancer. Overall and, where retained, classwise mean absolute SHAP summaries are shown. Attribution values quantify model contribution and do not establish biological causality.

**Figure S3G**

Multiclass LungCancer — class-specific panels 1/2

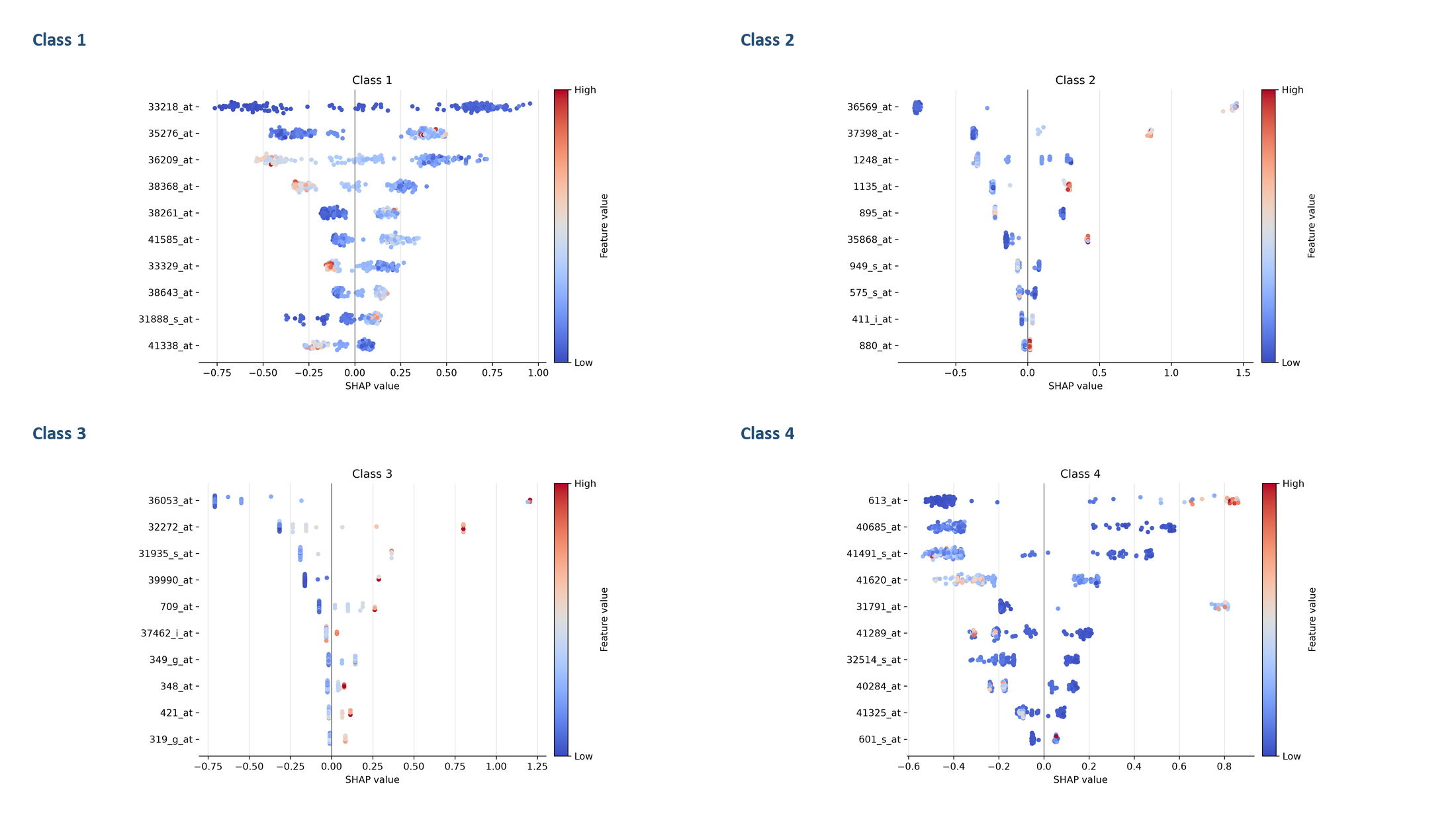

**Figure S3G (continued).** Class-specific SHAP distributions for Multiclass LungCancer. Each panel shows the magnitude and direction of feature contributions for the indicated class; red denotes higher feature values and blue denotes lower values.

**Figure S3G**

Multiclass LungCancer — class-specific panels 2/2

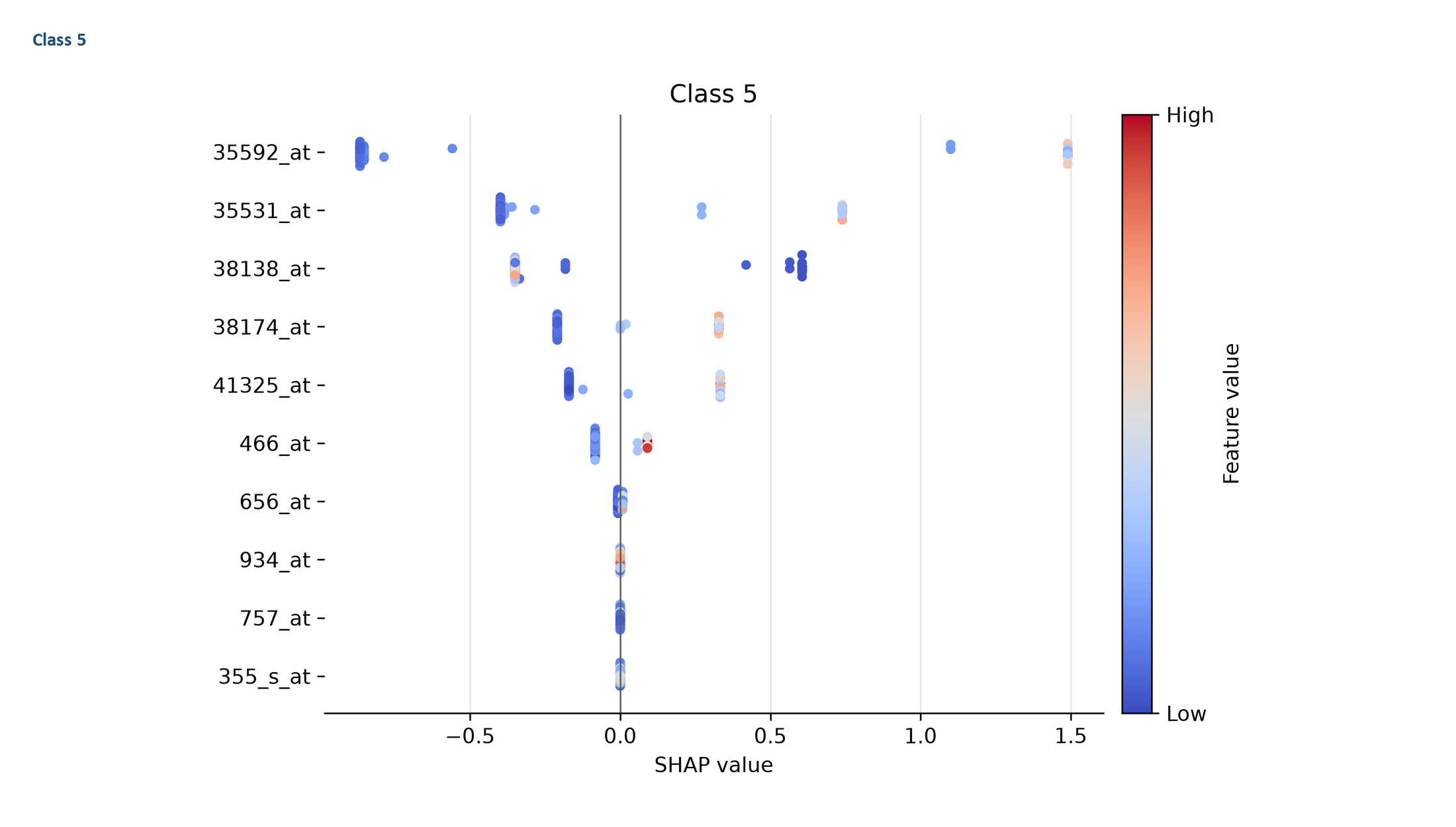

**Figure S3G (continued).** Class-specific SHAP distributions for Multiclass LungCancer. Each panel shows the magnitude and direction of feature contributions for the indicated class; red denotes higher feature values and blue denotes lower values.

**Figure S3H**

Multiclass Lymphoma — overview

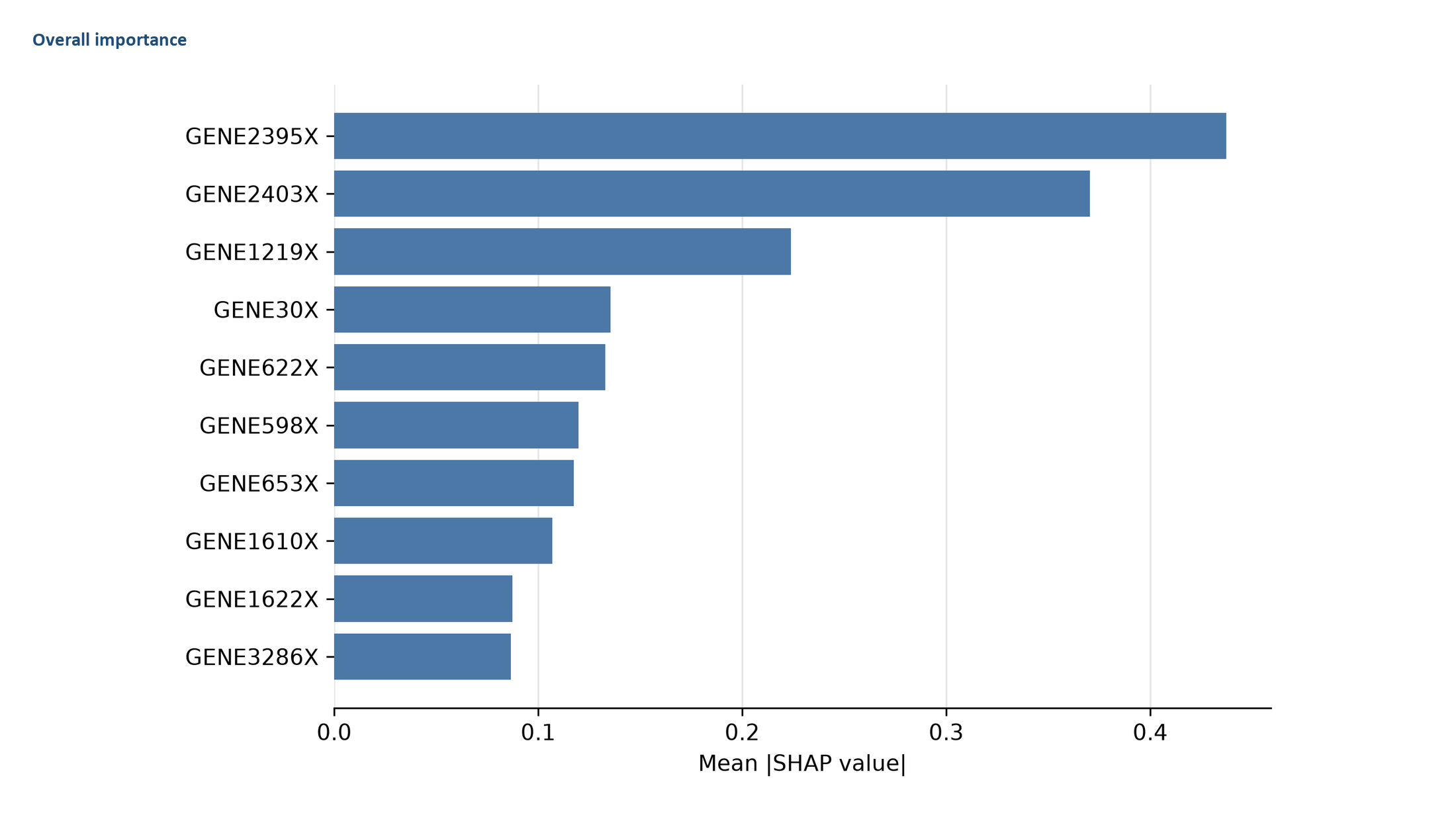

**Figure S3H.** Multiclass SHAP summary for Multiclass Lymphoma. Overall and, where retained, classwise mean absolute SHAP summaries are shown. Attribution values quantify model contribution and do not establish biological causality.

**Figure S3H**

Multiclass Lymphoma — class-specific panels 1/1

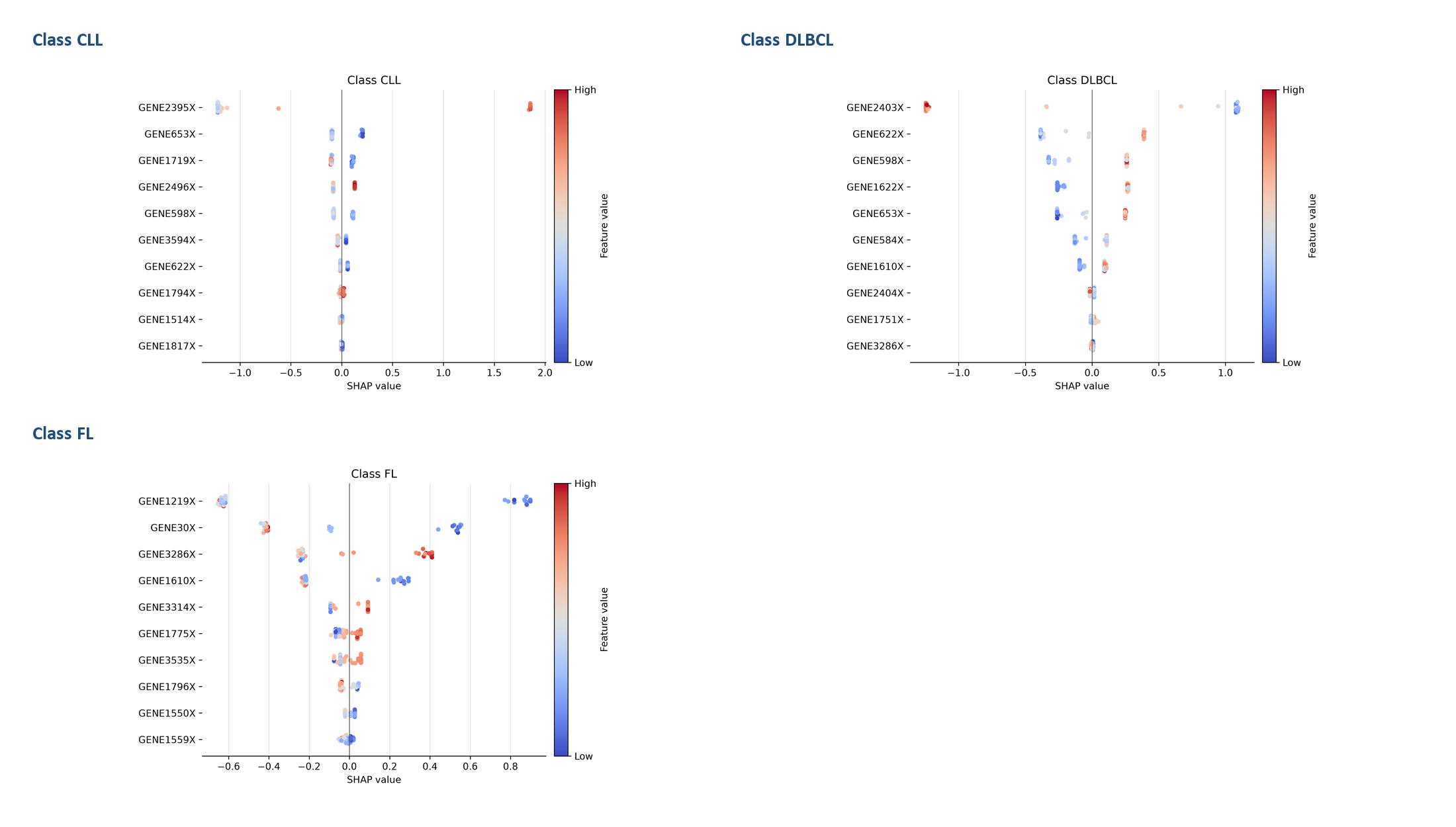

**Figure S3H (continued).** Class-specific SHAP distributions for Multiclass Lymphoma. Each panel shows the magnitude and direction of feature contributions for the indicated class; red denotes higher feature values and blue denotes lower values.

**Figure S3I**

Multiclass MLL — overview

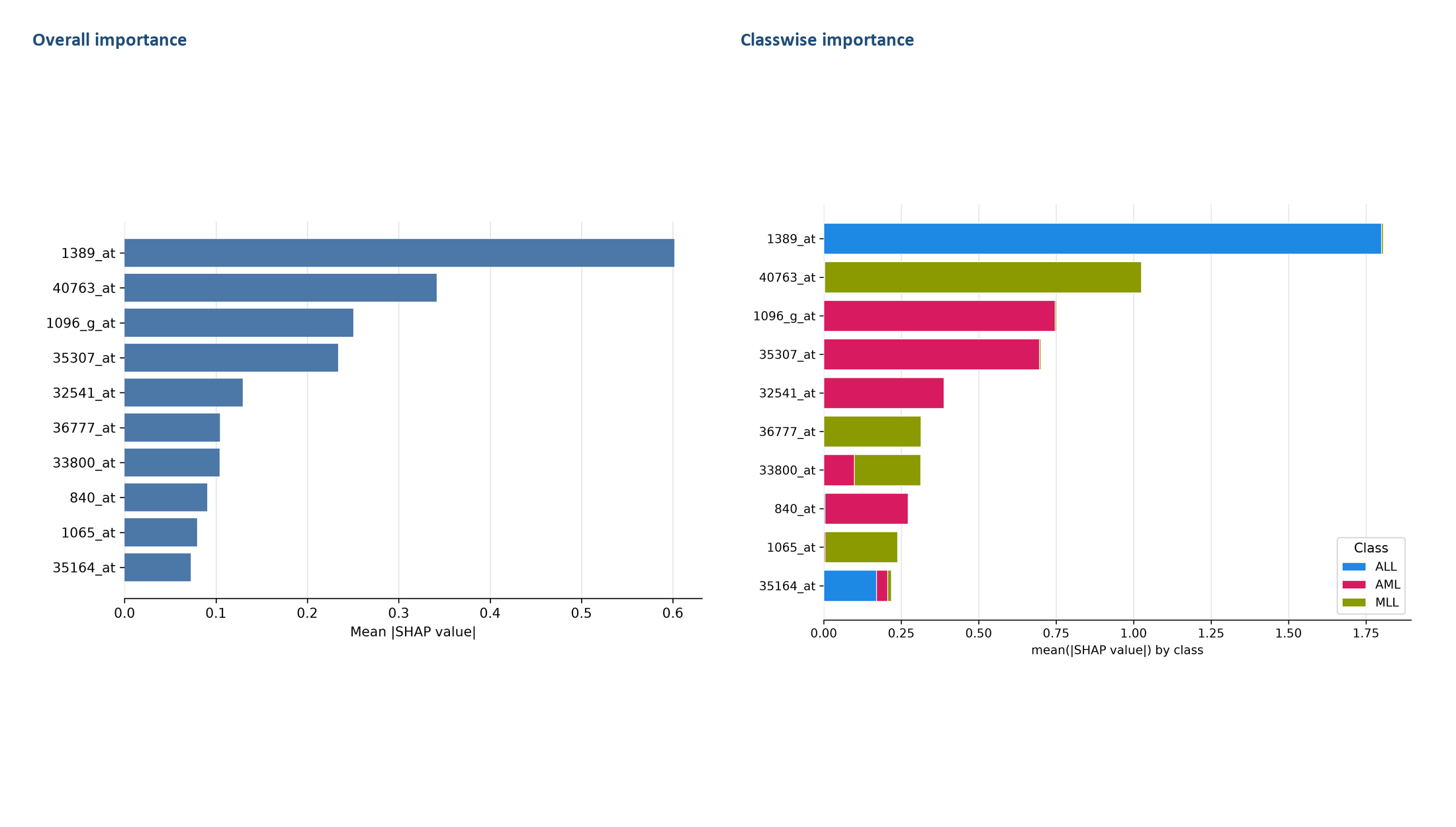

**Figure S3I.** Multiclass SHAP summary for Multiclass MLL. Overall and, where retained, classwise mean absolute SHAP summaries are shown. Attribution values quantify model contribution and do not establish biological causality.

**Figure S3I**

Multiclass MLL — class-specific panels 1/1

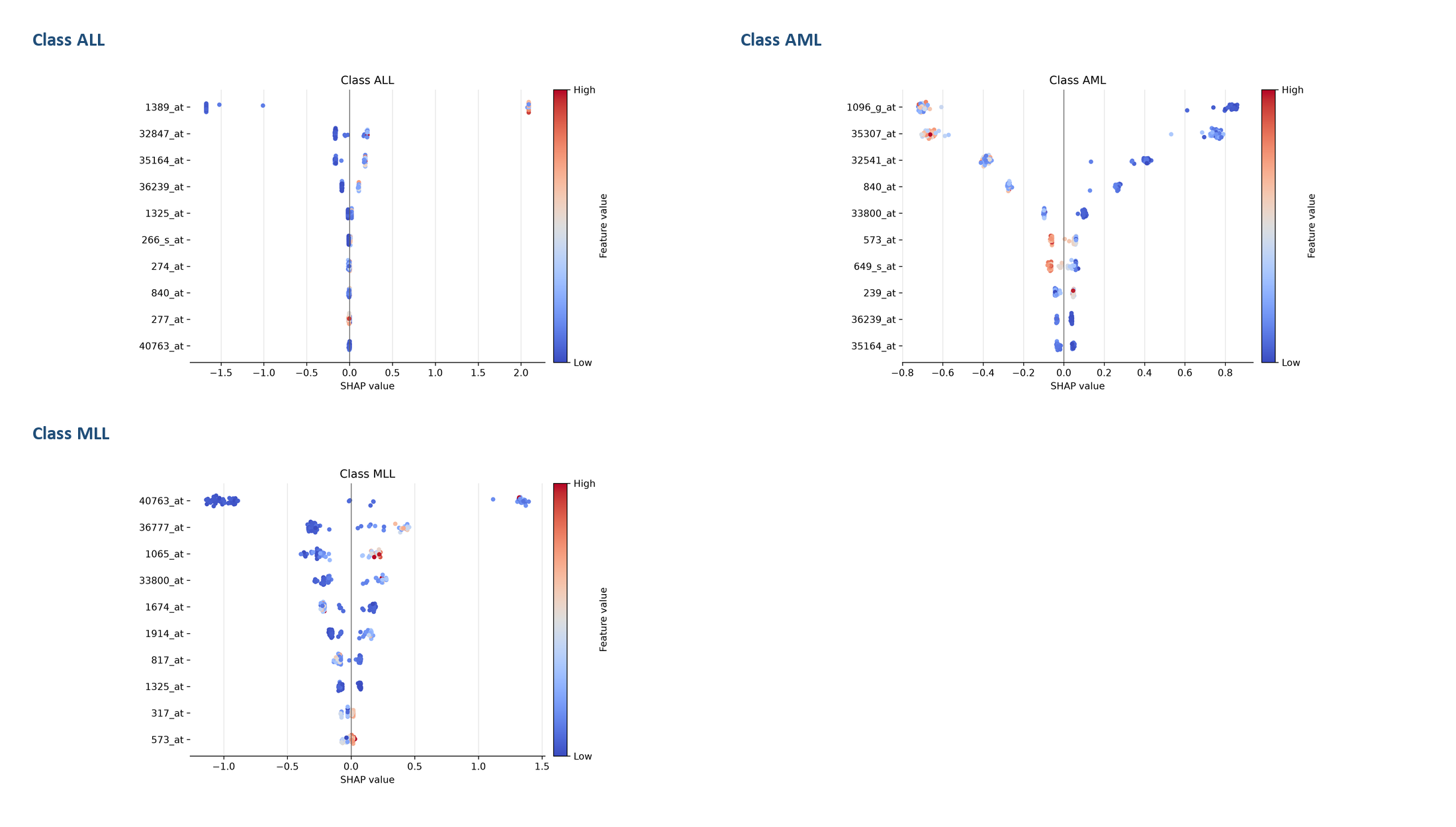

**Figure S3I (continued).** Class-specific SHAP distributions for Multiclass MLL. Each panel shows the magnitude and direction of feature contributions for the indicated class; red denotes higher feature values and blue denotes lower values.

**Figure S3J**

Multiclass SRBCT — overview

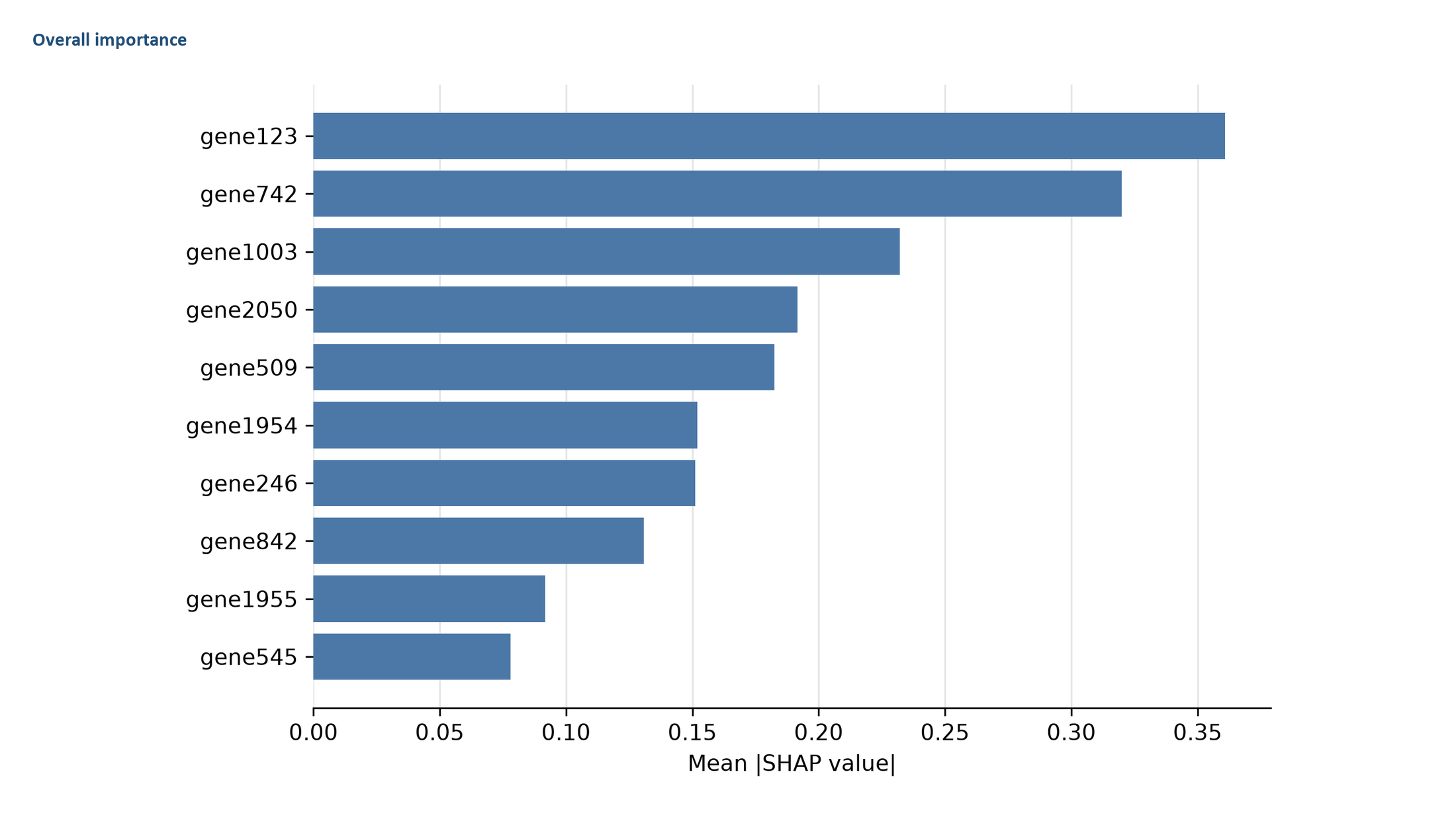

**Figure S3J.** Multiclass SHAP summary for Multiclass SRBCT. Overall and, where retained, classwise mean absolute SHAP summaries are shown. Attribution values quantify model contribution and do not establish biological causality.

**Figure S3J**

Multiclass SRBCT — class-specific panels 1/1

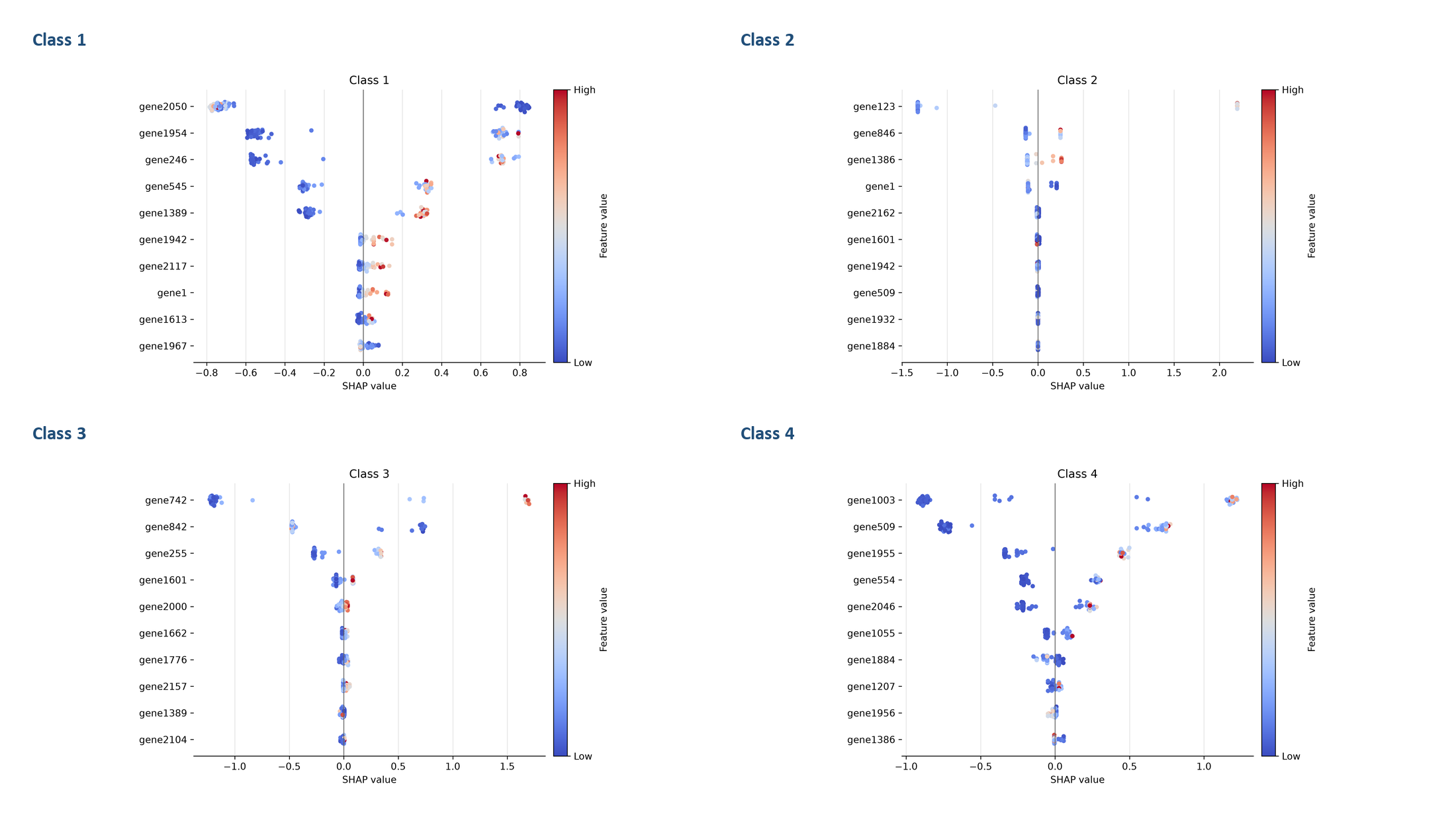

**Figure S3J (continued).** Class-specific SHAP distributions for Multiclass SRBCT. Each panel shows the magnitude and direction of feature contributions for the indicated class; red denotes higher feature values and blue denotes lower values.
